# Low-rank tensor decompositions reveal coupled spatiotemporal patterns of CSF tracer transport in the human brain

**DOI:** 10.64898/2026.09.14.750103

**Authors:** Andreas Solheim, Geir Ringstad, Per Kristian Eide, Marie E. Rognes, Evrim Acar, Kent-Andre Mardal

## Abstract

Intrathecal contrast-enhanced longitudinal MRI, or glymphatic MRI (gMRI), provides a unique clinical window into human cerebrospinal fluid (CSF) transport, interstitial fluid (ISF) interaction, and the glymphatic system. However, standard analyses flatten these multi-subject longitudinal data into univariate comparisons across regions of interest, destroying the underlying multi-way structure and obscuring coupled spatiotemporal dynamics. Here we analyze gMRI data from 92 patients (43 diagnosed with idiopathic normal pressure hydrocephalus [iNPH] and 49 reference subjects) using unsupervised non-negative low-rank tensor decompositions that treat tracer signal in CSF and brain parenchyma simultaneously. The analysis is stratified by sex to avoid confounding diagnosis with the sex imbalance between cohorts. We recover four replicable components, three of which resolve distinct transport pathways capturing CSF–ISF interaction: distribution of tracer in the supratentorial subarachnoid space and cerebral gray and white matter at 24 hours; transient early influx in regions consistent with transport along major cerebral arteries; and tracer influx to ventricular CSF, also called ventricular reflux. The first two components are positively correlated and thus establish a direct quantitative link between early-stage influx and 24-hour tracer distribution. Moreover, the expression of the two influx pathways varies significantly between REF and iNPH cohorts, demonstrating how unsupervised tensor decompositions provide an automated, data-driven framework to stratify patient groups and derive quantitative markers of solute transport in the human brain.

## 1 Introduction

Understanding solute transport via the cerebrospinal fluid (CSF) and interstitial fluid (ISF) is critical for deciphering the mechanisms underlying human brain waste clearance and neurological health (Rasmussen et al., 2022). The discovery of the glymphatic system (J. Iliff et al., 2012) and the re-discovery of dural meningeal lymphatic vessels (Aspelund et al., 2015; Louveau et al., 2015) have placed brain fluid transport at the center of studies of neurodegenerative disease and brain function. The key discovery of the glymphatic system is that CSF in the subarachnoid space (SAS) is directly coupled to ISF in the parenchyma, with perivascular spaces acting as the conduit. Recent studies using intrathecal contrast agents have also shown that perivascular spaces play a key role in molecular transport in humans (Eide & Ringstad, 2024; Yamamoto et al., 2024). However, while the coupled enrichment across CSF and parenchyma has been observed in humans (Eide & Ringstad, 2015; Ringstad et al., 2018; Wåhlin et al., 2026), the coupling itself has not been quantified. Doing so requires methods that treat CSF and parenchymal signal jointly, without collapsing the structure of multi-subject spatiotemporal data.

Quantifying these dynamics in humans relies on a rapidly expanding neuroimaging toolkit, aiming to capture brain clearance across a range of temporal and spatial scales. Recent advancements include contrast-enhanced *T*_1_-weighted MRI, where contrast is delivered either intrathecally (often referred to as glymphatic MRI, gMRI) (Eide & Ringstad, 2015; J. J. Iliff et al., 2013; Ringstad et al., 2018; Watts et al., 2019) or intravenously (Richmond et al., 2023; Wåhlin et al., 2026) to trace solute transport, alongside other non-invasive protocols measuring key quantities related to transport in brain tissue and CSF such as deformation, pulsation, SAS protein characterization, and dynamic diffusivity (Goa et al., 2026; Hirschler et al., 2025; Kiviniemi et al., 2016; Nwotchouang et al., 2021; Storås et al., 2026; Wen et al., 2025).

Each of these advancements represents dynamic imaging techniques that generate complex, multi-subject, multi-dimensional longitudinal datasets spanning timeframes from milliseconds to several days. From a dimensional point of view, this type of data is 5D, including three spatial dimensions for each time point and each subject. Conventional data analysis approaches remain largely dependent on univariate comparisons across pre-defined regions of interest (ROIs). However, flattening multidimensional data arrays into matrices eliminates the underlying multi-way structure of the data and obscures coupled spatial, temporal or subject interactions, making it more difficult to characterize population-level transport dynamics across heterogeneous patient cohorts.

Low-rank tensor decompositions offer an unsupervised, data-driven framework aimed directly at these analytical challenges. The core element is to keep the representation of longitudinal, multi-subject neuroimages as tensor-valued data. Techniques such as the non-negative canonical polyadic (CP) decomposition can then decouple complex data patterns into separate spatial, temporal, and subject modes, without *a priori* spatial assumptions (Ballard & Kolda, 2025; Kolda & Bader, 2009). While tensor decompositions are widely established in functional neuroimaging (EEG/fMRI) (Acar et al., 2007; Beckmann & Smith, 2005; Cong et al., 2015; De Vos et al., 2007; Erol & Hunyadi, 2022; Martınez-Montes et al., 2004; Miwakeichi et al., 2004; Mørup et al., 2008; Mørup et al., 2006, 2025; Williams et al., 2018) and broader data science (Acar & Yener, 2009; Ballard & Kolda, 2025; Bro, 1997; Sidiropoulos et al., 2017; Smilde et al., 2004), their application to CSF–ISF exchange, solute transport, and brain fluid clearance in the human brain remains unexplored.

Here, we study solute transport in the human brain using a gMRI dataset of 92 subjects recruited while under investigation for various CSF disorders at Oslo University Hospital. The dataset includes 43 patients diagnosed with idiopathic normal pressure hydrocephalus (iNPH) and 49 reference subjects (REF) with no CSF disorder or neurological disease following clinical work-up. All subjects underwent intrathecal injection of gadobutrol, with MRI performed prior to injection and approximately 2, 4, 8, and 24 hours after injection (Ringstad et al., 2017, 2018). Brain solute transport and CSF flow patterns are known to vary substantially both between individuals and across neurological conditions (Eide, Valnes, et al., 2021; Ringstad et al., 2018), making population-level patterns difficult to characterize from individual comparisons. By simultaneously treating tracer signal in the CSF and parenchyma in a cohort of patients, we investigate to what extent unsupervised learning, in the form of low-rank tensor decompositions, can discover, quantify, and differentiate composite patterns of tracer dynamics in the brain.

Using the non-negative CP decomposition, we represent the gMRI data as a set of distinct patterns, termed components, that decouple tracer dynamics into separate spatial, temporal, and subject modes, and we establish their robustness through repeated split-half cross-validation (Adali et al., 2022). Jointly evaluating the three modes allows each component to be interpreted in terms of physiological transport features and to compare its expression between cohorts. Among the four patterns we identify, one component directly quantifies regions involved in tracer distribution in the supratentorial SAS and cerebrum 24 hours after injection, while a second identifies transient influx consistent with early tracer transport along major cerebral arteries. The positive correlation between the two links early-stage influx to 24-hour tracer distribution. A third component captures ventricular reflux, a pattern characterized by tracer entering ventricular CSF and previously graded by expert raters (Eide et al., 2020), which is of particular clinical interest for classifying shunt-responsive iNPH patients. Its expression is significantly higher in iNPH than in REF subjects, correlates positively with age, and provides a continuous, data-driven alternative to rater-based grading. Each component illustrates areas in the CSF and parenchyma that evolve over the same time profile, allowing us to quantitatively describe the CSF–ISF coupling central to the glymphatic framework.

## 2 Methods and materials

### 2.1 Patients and MRI protocol

#### 2.1.1 Patients and approvals

Subjects in this study were recruited while under clinical work-up for suspected CSF disorders, with imaging performed at the Intervention Centre, Oslo University Hospital–Rikshospitalet. The dataset includes a total of 92 subjects who underwent a gMRI protocol, described in detail in the following section (Section 2.1.2). Among the subjects included in this study, 43 (28 male, 15 female) were diagnosed with idiopathic normal pressure hydrocephalus (iNPH), a condition characterized by gait disturbance, urinary incontinence, and dementia, with histopathological overlap towards Alzheimer’s disease (Eide, 2025; Malm & Eklund, 2006). A hallmark of iNPH is altered CSF dynamics, historically defined through diverse clinical criteria, including hyperdynamic aqueductal flow imaged by phase-contrast MRI (Bradley Jr, 2015), increased CSF outflow resistance measured during infusion testing (Jacobsson et al., 2018), and elevated intracranial pressure wave amplitudes (Wagshul et al., 2011). The remaining 49 patients (11 male, 38 female) were not diagnosed with any CSF disorder or neurological disease following clinical work-up and serve as a reference group (REF), representing the closest available approximation of healthy controls. Subject demographics, including age, height, and weight, across groups are summarized in Table 1.

**Table 1:** Characteristic data for patients in the dataset. We show the median and inter-quartile range of age, height and weight for the entire dataset, as well as when splitting the dataset by diagnosis (REF, iNPH) and sex.

| Group | Subjects | Age | Height [cm] | Weight [kg] |
| --- | --- | --- | --- | --- |
| REF | 49 (11 Male/ 38 Female) | 37 [30, 46] | 170 [166, 176] | 80 [70, 90] |
| iNPH | 43 (28 Male/ 15 Female) | 69 [54, 73] | 176 [168, 180] | 86 [69, 95] |
| Male | 39 (11 REF/ 28 iNPH) | 65 [49, 73] | 180 [177, 186] | 90 [82, 100] |
| Female | 53 (38 REF/ 15 iNPH) | 38 [30, 53] | 168 [164, 170] | 73 [64, 86] |
| **Total** | **92 (49 REF/ 43 iNPH)** | **47 [32, 69]** | **173 [167, 180]** | **82 [70, 91]** |

Collection of data analyzed in this study was approved by the Regional Committee for Medical and Health Research Ethics (REK) of Health Region South-East, Norway (2015/96), the Institutional Review Board of Oslo University Hospital (2015/1868) and the National Medicines Agency (15/04922-7), and was conducted following the ethical standards of the Declaration of Helsinki of 1975 (revised in 1983). Study participants were included after written and oral informed consent. Parts of the dataset have been included in previous studies of human glymphatic function conducted at the University Hospital of Oslo (Ringstad et al., 2017, 2018) in the years 2015–2019. No new data were collected for the present work.

#### 2.1.2 Glymphatic MRI (gMRI) protocol

Patients included in this work underwent the gMRI protocol introduced in Ringstad *et al*. (Ringstad et al., 2017, 2018), involving longitudinal MRI of an intrathecal contrast enhancing agent aiming to illustrate brain-wide CSF transport and clearance. In each patient, a dose of 0.5 mmol (0.5 ml of 1.0 mmol/ml gadobutrol; Gadovist, Bayer Pharma AG) was administered intrathecally by an interventional neuroradiologist. MRI was performed prior to injection, as well as approximately 2, 4, 8, and 24 hours after injection. The observed signal changes reflect tracer distribution in CSF spaces and tissue compartments and should not be interpreted as a direct measurement of any single clearance mechanism.

In each patient, the longitudinal MRIs were aligned to the pre-injection image using FreeSurfer (Dale et al., 1999) (version 6) and resampled to the FreeSurfer-standard 256 *×* 256 *×* 256 grid with 1 mm resolution. We segment the pre-injection MRI using the Desikan–Killiany atlas through FreeSurfer (Dale et al., 1999; Desikan et al., 2006), providing a detailed segmentation of the brain parenchyma and ventricles. We then expand this segmentation to include the CSF in two steps: First, we use *T*_2_-weighted imaging to identify a CSF mask. We then associate each voxel in this mask with the closest label in the segmentation of the parenchyma. Accordingly, the final segmentation includes patient-specific regions of interest (ROIs) in both the brain parenchyma and CSF in the ventricles and the SAS. Figure 1 shows some of the key major regions of the brain we can identify from this segmentation.

**Figure 1:**
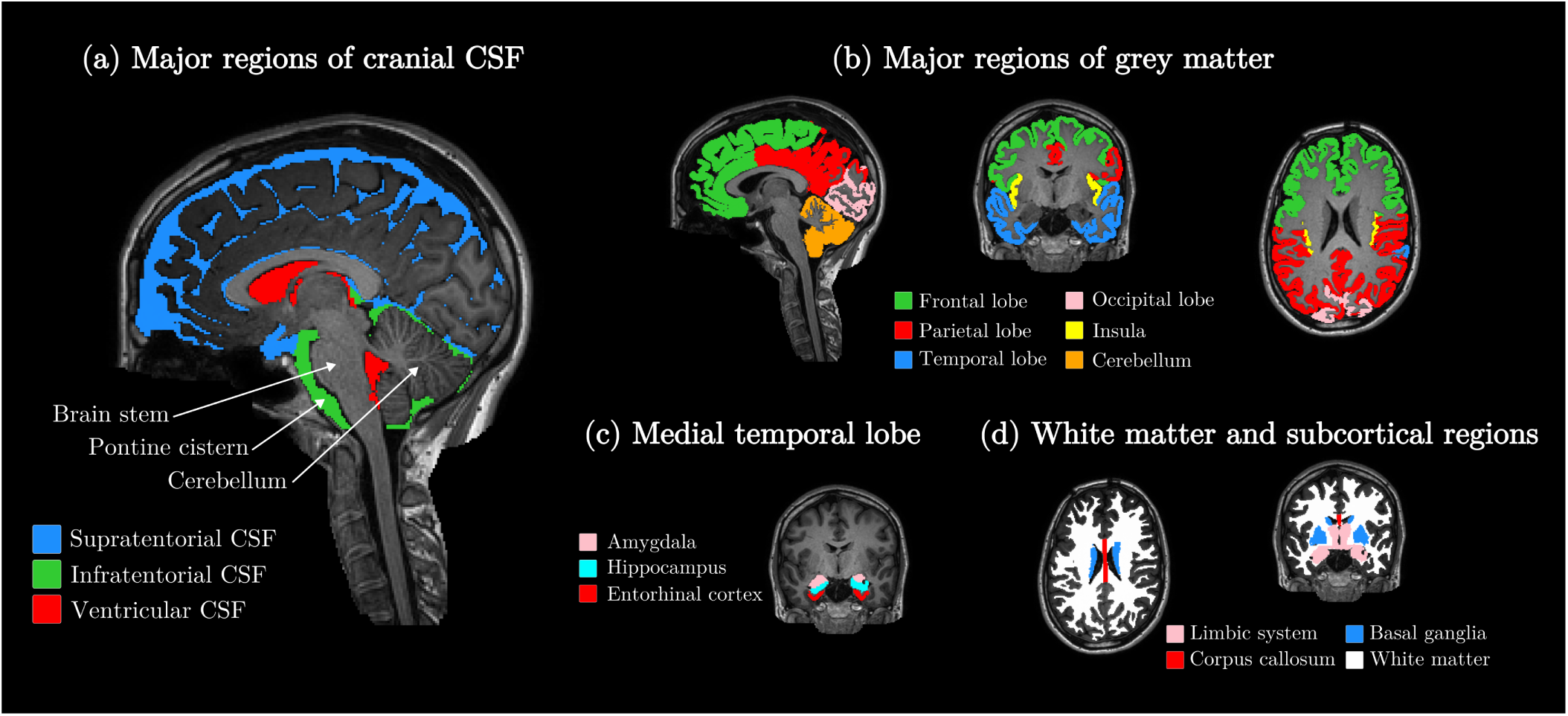
Overview of some of the major anatomical regions most relevant in this work. The illustration is built using the same segmentation as when computing the CP decomposition, and we use the same slices as in subsequent figures. We group some ROIs into larger regions in the interest of clarity: (a) Illustration of the major CSF regions we use in this work (b) gray matter regions split into the major brain lobes, as well as the insula and cerebellar gray matter (c) Main regions of the medial temporal lobe (d) White matter and some subcortical gray matter structures.

### 2.2 Tensor analysis of gMRI data

#### 2.2.1 Representing gMRI data as a tensor

When converting longitudinal image data into tensors we take advantage of the shared representation provided by the patient-specific segmentations. Because each ROI labels a specific region of the brain, we can directly compare the tracer transport between individuals. We measure the evolution of the MRI tracer in each subject by comparing the *T*_1_-weighted signal intensity at each of the four post-injection time-points with the pre-injection MRI. In each voxel, we divide the post-contrast signal by the pre-contrast signal and then find the median voxel value in each ROI of the corresponding patient segmentation. We only include regions which are present in all patients in the dataset, leaving a total of 245 included ROIs. By combining the median signal increase in each ROI, we obtain a vector for each subject. We then combine the vectors for each of the four time-points in each subject column-wise to construct a 245 *×* 4 matrix. Each subject matrix is then stacked to construct a third-order tensor of size 245 *×* 4 *× n*, where *n* is the number of included subjects. As such, the 5D tensor is represented as a 3D tensor in the following. Figure 2a illustrates how the data is agglomerated in ROIs to construct a vector, and how we proceed to construct the tensor.

**Figure 2:**
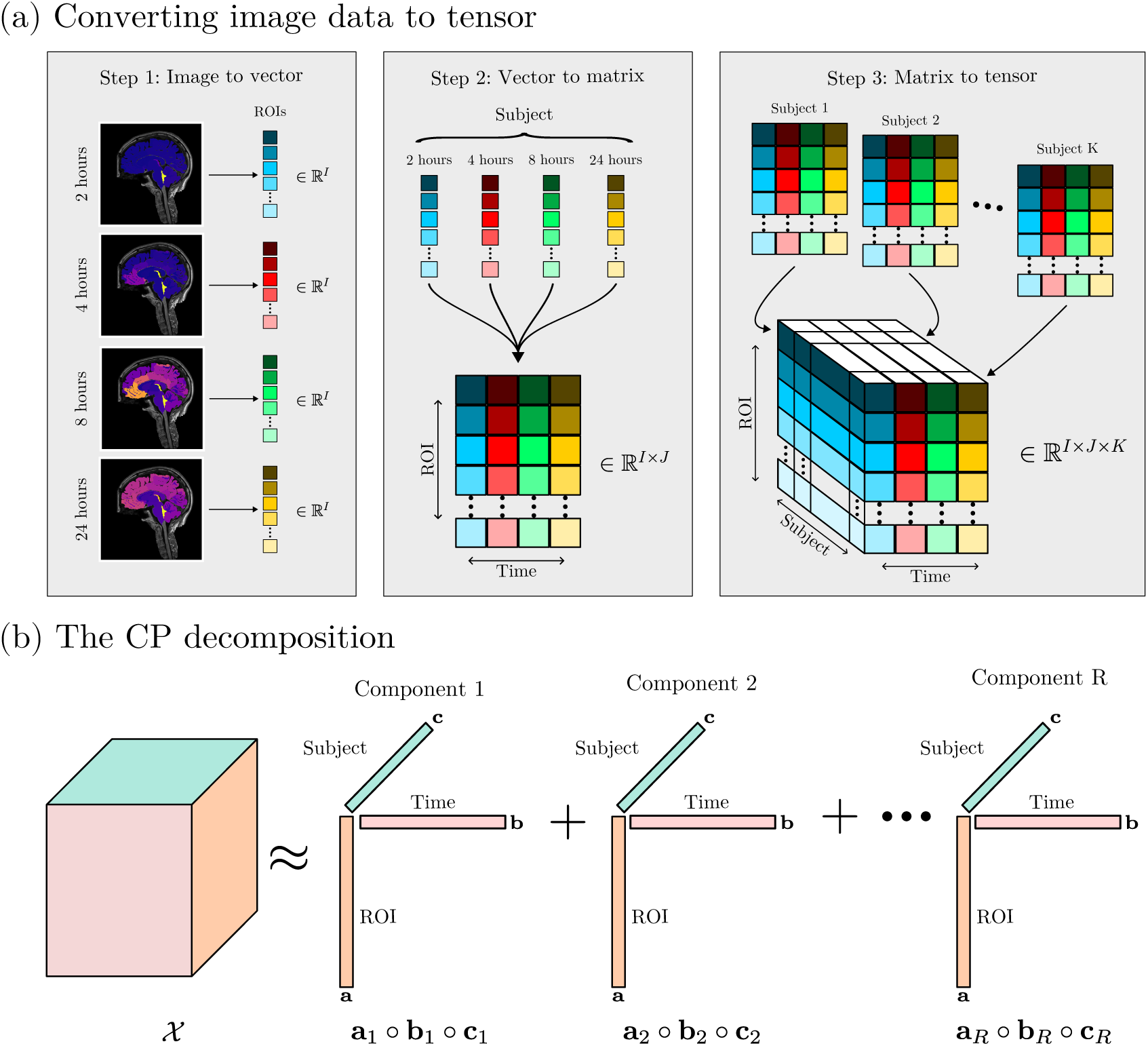
The CP decomposition of gMRI data. Figure (a) shows how multi-subject longitudinal image data is converted into a tensor. In Step 1, we compute the median signal increase in each ROI to create a vector for each subject at each time-point. In Step 2, we stack the vectors from each time-point column-wise to create a matrix. Finally, in Step 3 we stack the subject-matrices to create a third-order tensor. Figure (b) shows how the CP decomposition approximates the original tensor as a sum of rank-one tensors.

#### 2.2.2 The canonical polyadic (CP) decomposition

The CP decomposition, also known as CANDECOMP (Canonical Decomposition) or PARAFAC (Parallel factor analysis) (Carroll & Chang, 1970; Harshman, 1970; Hitchcock, 1927; Möcks, 1988), is a low-rank tensor decomposition technique that gives a unique factorization (up to scaling and permutation) under mild conditions (Kruskal, 1977). Because the relative signal compared to baseline is non-negative, we apply this as an additional constraint and consider the non-negative CP decomposition. We compute CP using the non_negative_parafac method implemented in TensorLy (Kossaifi et al., 2019) (v.0.9.0).

Given a third order tensor *X ∈* R*^I×J×K^* the goal of the non-negative CP decomposition is to approximate *X* as the sum of *R* rank one tensors, with non-negative coefficients, for a given integer *R* ⩾ 1. Specifically, we aim to find triplets of vectors 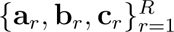, with 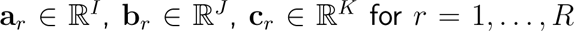, and corresponding scaling factors *λ_r_* ⩾ 0, which minimize:

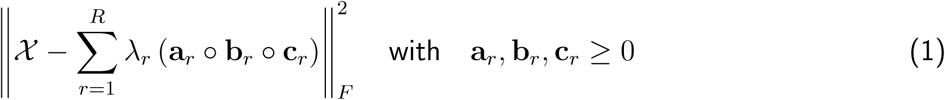

where *||·||_F_* denotes the Frobenius norm and *◦* denotes the outer product. The factor *λ_r_*is computed such that *||***a_r_**||_2_ = ||**b_r_** ||**c_r_**||_2_ = 1, where *||·||*_2_ denotes the *ℓ*^2^-norm. In element-wise scalar form, the modeled tracer signal increase in ROI *i*, at post-injection time-point *j*, for subject *k* is then approximated as:

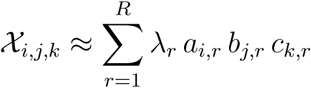

In the following we refer to each triplet {**a***_r_,* **b***_r_,* **c***_r_*} as a *component*, and we refer to the triplet with index *r* as the *r*th component. The value *λ_r_* quantifies the relative importance of each component when reconstructing the data. When computing a decomposition with *R* total components we refer to the decomposition as an *R*-component model.

For the gMRI dataset, *I* = 245, *J* = 4, and *K* = *n*, with *n* being the number of included patients, for a given component *r*, the vector **a***_r_* encodes spatial, ROI-specific information, **b***_r_* encodes time-specific information while **c***_r_* encodes patient-specific information. We refer to each of these three axes of the tensor as *modes*; i.e., ROI/spatial mode, time mode, and subject mode, respectively. Figure 2b shows how the modes and components relate to the original tensor.

#### 2.2.3 Variance-normalization and interpretation of CP components

Tracer data from gMRI display significant scaling differences between CSF and parenchyma due to the different extra-cellular volume fraction, and some ROIs will have large variations in tracer enrichment across the four time points. In order to mitigate the effect of certain regions and time points dominating the end outcome, we divide the value in each ROI by the standard deviation across all time-points and all included subjects before computing the CP decomposition. Accordingly, the ROI mode of each component shows the variance-normalized importance of each region compared to others. The spatial modes should therefore not be interpreted directly as reflecting tracer enrichment, but rather the relative involvement of each ROI. ROIs with lower overall tracer accumulation may therefore exhibit prominent spatial component weights if their temporal dynamic shifts are marked relative to their baseline variance.

#### 2.2.4 CP model selection

While the CP decomposition of a tensor is unique, the objective function in (1) is potentially highly non-convex and must be solved through iterative methods. The models may therefore be sensitive to the initialization of the algorithm. We therefore use a strategy of repeated randomized initializations, aiming to reduce the risk that the models we analyze have converged to a local minimum. The choice of the best model is based on two criteria: First, the model must achieve a low error as defined by (1) and second, it must be consistent with other models with similar error. We measure consistency using the Factor Match Score (FMS), which evaluates the similarity of two decompositions {**a**_1_ … **a***_R_*}, {**b**_1_ … **b***_R_*}, {**c**_1_ … **c***_R_*} and {**â**_1_ … **â***_R_*}, {**b̂**_1_ … **b̂***_R_*}, {**ĉ**_1_ … **ĉ***_R_*}:

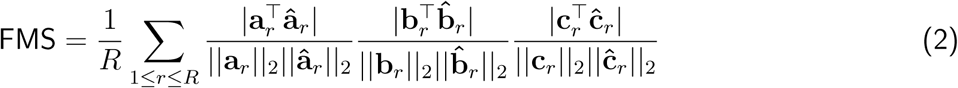

The dual model selection criterion (low error and consistent models) ensures that the model has converged to a minimum that can be reached consistently, increasing the likelihood that it reflects a global minimum of (1).

When computing a CP model, choosing the correct number of components *R* is a critical hyperparameter. Unlike matrix-based methods, computing the rank of a third-order tensor is NP-hard (Kolda & Bader, 2009), meaning that *R* is a hyperparameter that must be tuned through an iterative approach. In this work, we focus on choosing *R* such that the patterns we compute are *replicable* (Adali et al., 2022). The goal is to identify patterns which would also be present in a different set of patients belonging to the same diagnosis groups and following the same MRI protocol but collected independently from the present dataset. We estimate the replicability of a *R*-component CP model using repeated split-half cross validation: we repeatedly split the dataset into non-overlapping halves and then compute the CP model on each half on the dataset. The resulting ROI and time modes from the two models are then compared using FMS. By repeating this 100 times we obtain an estimate of how sensitive a given *R*-component model is to the sampling of the dataset. We consider models with an average FMS score greater than 0.9 to be replicable.

## 3 Results

Prior to interpreting the components we compute using CP we perform a replicability analysis using 100 times repeated split-half cross-validation and compare models using the Factor Match Score (FMS), described in Section 2.2.4. Figure 3 shows the distribution of FMS scores when stratifying the dataset by diagnosis (REF, iNPH) or sex (Male, Female) for CP models with *R* between 2 and 6. The average FMS score for each model is shown in Table 2 and we find that in most dataset subsets (Female, REF, iNPH), 4 is the largest number of components that yield a replicable model, except in the male subset where 3 components gives the largest replicable model.

**Figure 3:**
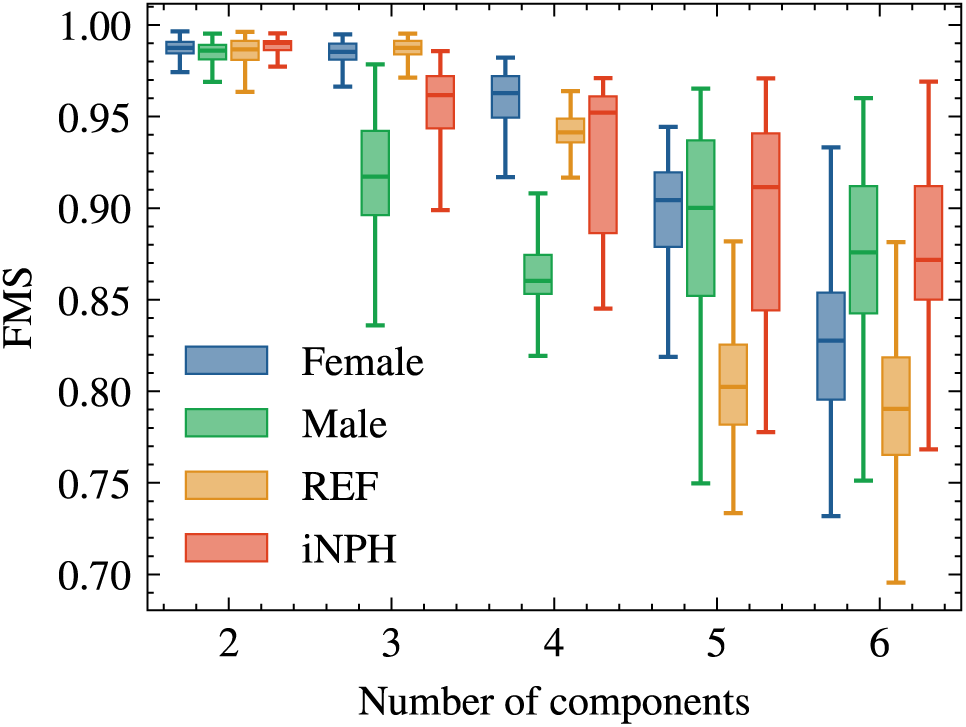
Distribution of FMS scores when splitting the dataset into subsets based on sex or diagnosis, as a function of the number of CP components included in the model. We split each data subset randomly into two non-overlapping groups and compute the CP decomposition for each half and compute the similarity of the time and ROI modes of the computed decompositions using FMS. This procedure is repeated 100 times for each data subset and we report the distribution.

**Table 2:** Average FMS score of the distributions shown in Figure 3. We consider CP models with an average FMS score of 0.9 or greater as being replicable. These instances are emphasized in bold.

| Group | Subjects | $R = 2$ | $R = 3$ | $R = 4$ | $R = 5$ | $R = 6$ |
| --- | --- | --- | --- | --- | --- | --- |
| Male | 39 | **0.99** | **0.92** | 0.88 | 0.89 | 0.87 |
| Female | 53 | **0.99** | **0.98** | **0.96** | 0.89 | 0.83 |
| REF | 49 | **0.98** | **0.99** | **0.94** | 0.81 | 0.79 |
| iNPH | 43 | **0.99** | **0.95** | **0.93** | 0.90 | 0.88 |

In our analysis, we consider the dataset stratified by sex or diagnosis due to the significant population skew among patients in the dataset. Table 1 shows that while the REF group is predominantly female (11 male, 38 female), the iNPH group is predominantly male (28 male, 15 female). Although there is currently no indication that sex is a primary determinant of tracer transport in the human brain, this imbalance in the dataset potentially makes it difficult to distinguish diagnosis-related effects from sex-related ones. In the following, we focus our analysis on the female subset in particular, because it is the largest subset of patients in this dataset. Other ways of splitting the dataset (Male, REF, iNPH) yield generally consistent patterns. In the Supplementary Material we provide a condensed analysis of patterns arising from other ways of stratifying the dataset.

### 3.1 Analysis of the 4-component CP model of the female gMRI data

On the female subset, we identify four components (the *static*, *24-hour supratentorial*, *early-influx*, and *ventricular reflux*) that express distinct tracer transport patterns, which we name according to their key characteristic features (Figure 4). We order the components in the order of their relative importance *λ_r_* to reconstruct the variance-normalized data. The static component has a constant time mode and isolates ROIs where tracer enrichment shows low variation over the measured period. This component corresponds closely to the ROI-wise average over all time points and subjects and is not interpreted as a transport pathway. The 24-hour supratentorial component rises to a peak at 24 hours and couples supratentorial SAS with cerebral gray and white matter, indicating direct CSF–ISF exchange. The early-influx component time profile peaks at 8 hours and returns to zero by 24 hours, with high spatial weights in the infratentorial SAS, prefrontal cortex, insula, and medial temporal lobes; its expression is significantly higher in REF subjects. The ventricular reflux component shares the 8-hour peak in the time profile but remains elevated at 24 hours, with spatial weights dominated by ventricular CSF and periventricular white matter; its expression is significantly higher in iNPH subjects. Table 3 lists some of the key features and relative weights of each component. In Figure 4 we plot each of the four components, showing time mode (left column), subject mode (second left column) and spatial mode (right and second right columns) together, aiming to illustrate any identifiable patterns. The spatial mode is also further illustrated in Figure 5, which shows a sagittal, coronal and axial slice of the four components. This figure is also complemented by Figure 1 which labels relevant structures in the corresponding slices. Each component is described in detail below.

**Figure 4:**
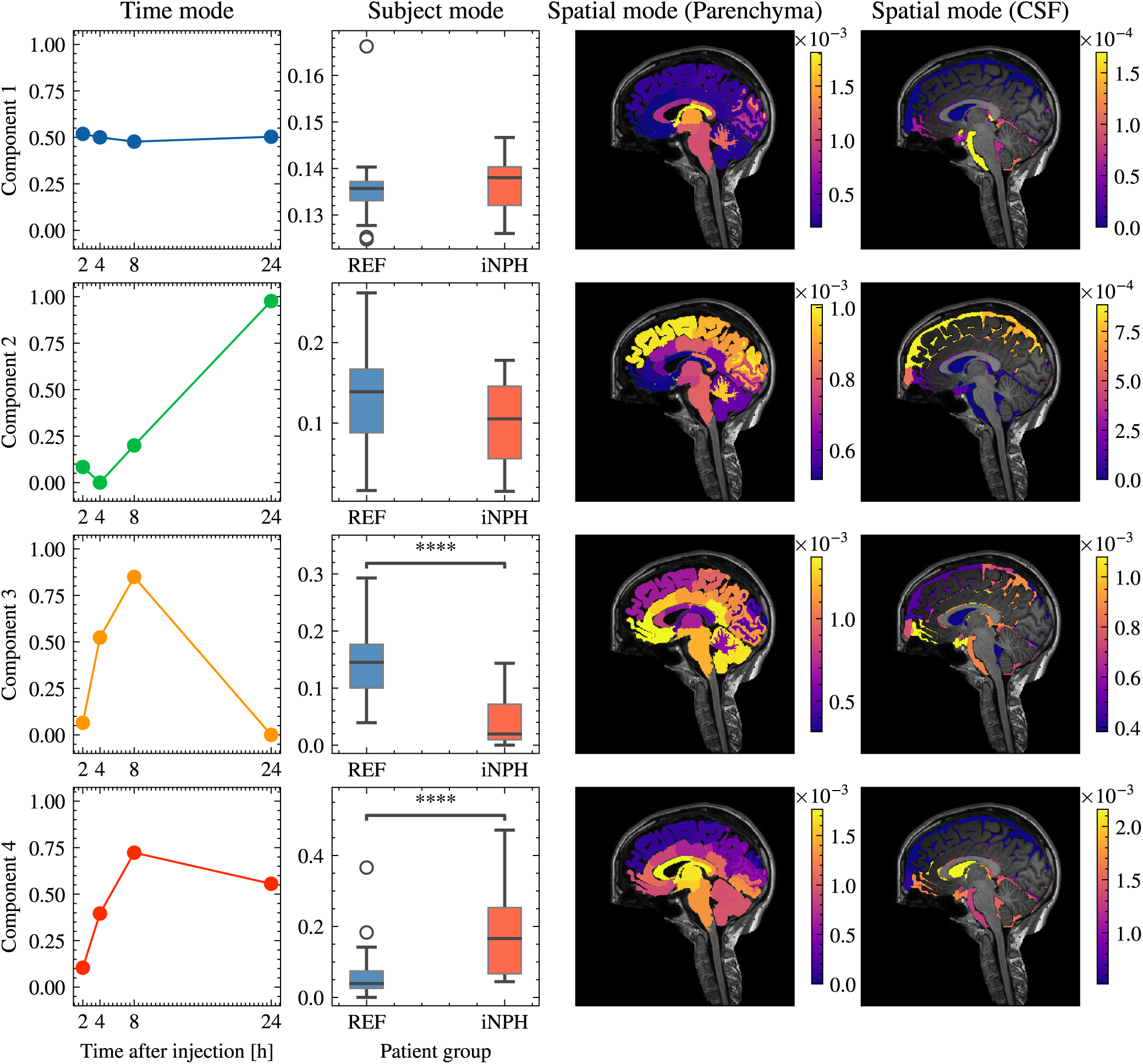
Four-component non-negative CP decomposition of the female subset. Components 1–4 correspond to the static, 24-hour supratentorial, early-influx, and ventricular reflux components respectively. Columns from left to right show the time mode, subject mode (REF vs. iNPH), and a sagittal view of the spatial mode in the parenchyma and CSF (see also Figure 5). The static component has a constant time mode and equal expression across groups. The 24-hour supratentorial component increases to a peak at 24 hours, and shows no significant group difference. The early-influx and ventricular reflux components share a peak at 8 hours but differ at 24 hours, where the early-influx coefficient returns to zero while ventricular reflux remains elevated. Their subject modes separate the cohorts in opposite directions, with early-influx expressed more strongly in REF and ventricular reflux in iNPH. Mann–Whitney U test, *∗ ∗ ∗∗ p <* 0.0001.

**Figure 5:**
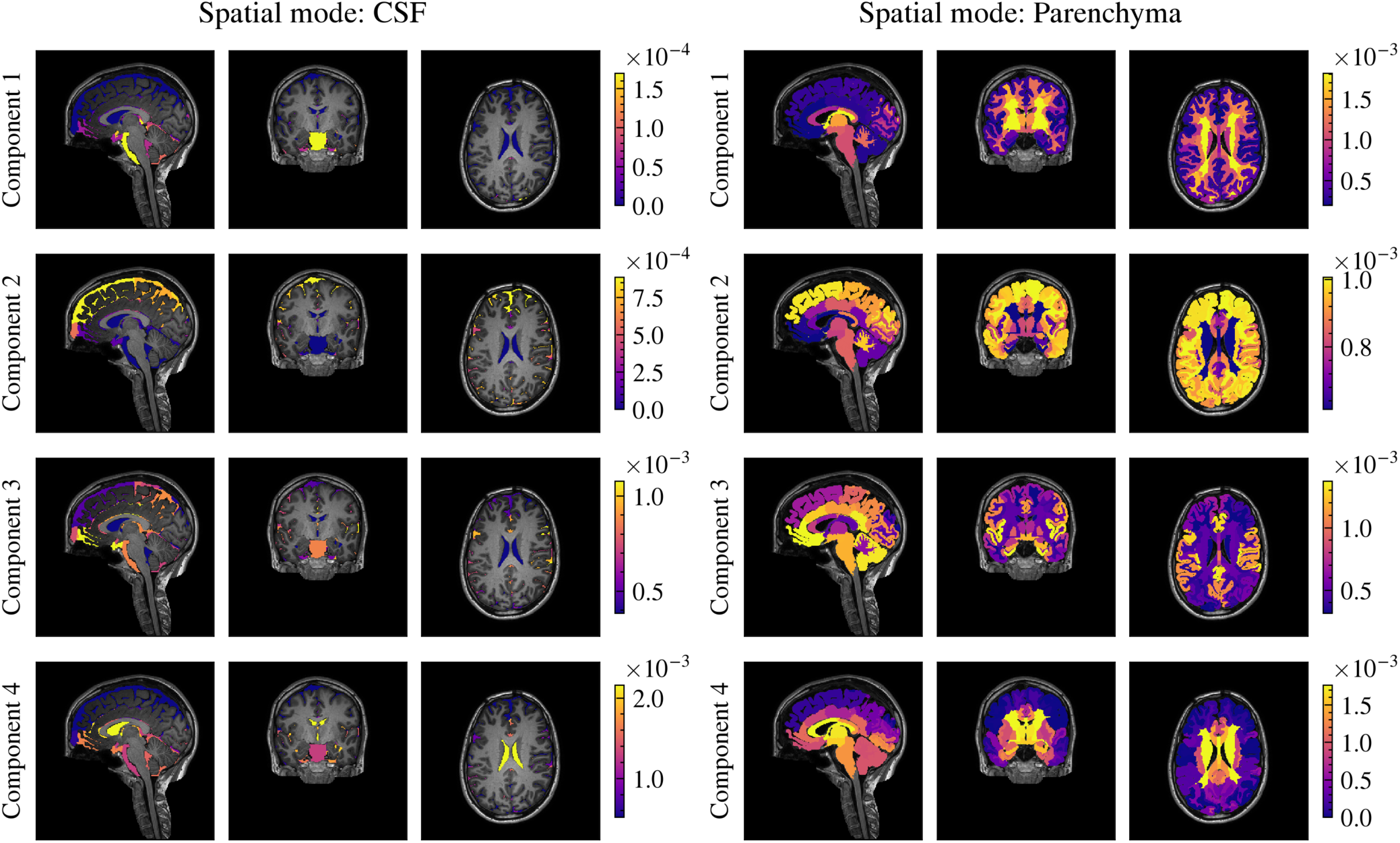
Sagittal, coronal, and axial slices of the 3D spatial modes for CSF (left panel) and parenchyma (right panel), corresponding to Figure 4. Rows are components 1–4 as in Figure 4; anatomical structures in these slices are labeled in Figure 1. In the CSF, the static component isolates the infratentorial CSF and particularly the pontine cistern; the 24-hour supratentorial component has elevated coefficients in superior regions of the supratentorial SAS; the early-influx component involves the infratentorial SAS and supratentorial SAS near the frontal lobe; and the ventricular reflux component is dominated by ventricular CSF. In the parenchyma, the static component emphasizes deep white matter, the 24-hour supratentorial component cerebral gray and white matter, the early-influx component prefrontal cortex, insula, and medial temporal lobes (particularly entorhinal cortex), and the ventricular reflux component periventricular white matter, basal ganglia, and parts of the medial temporal lobes.

**Table 3:** Summary of the 4-component CP model on the female subset. We compute the relative importance of each component *λ̃_r_*, scaling by the largest *λ_r_* (which is *λ*_1_) and summarize the main features of each component. The static component corresponds to the ROI-wise average of all time points and subjects in the dataset (after scaling by the standard deviation) and therefore has the highest weight *λ*. The value *λ̃_r_* of the other components should be interpreted as the relative importance of a given component compared to the static component.

| Component | Name | $\tilde{\lambda}_r = \lambda_r / \lambda_{max}$ | Main features |
| --- | --- | --- | --- |
| Component 1 | Static | 1 | Regions where the tracer concentration varies little over time (consistently high or low concentration). Time-average due to scaling. |
| Component 2 | 24-hour supratentorial | 0.22 | Tracer distribution at 24 hours, mainly in supratentorial SAS and cerebrum. |
| Component 3 | Early-influx | 0.16 | Transient early-stage tracer transport from the infratentorial SAS, peaking at 8 hours and clearing by 24.<br>Higher expression among REF subjects. |
| Component 4 | Ventricular reflux | 0.11 | Tracer entering ventricular CSF, peaking at 8 hours but remaining elevated at 24.<br>Higher expression among iNPH subjects. |

#### 3.1.1 Component 1: The static component

The static component (component 1, first row Figure 4) shows an essentially constant coefficient at all time points and no significant difference between patient groups. The equal weighting of all time points and subjects means this component is essentially an average, which is why it has the highest relative importance (Table 3). Its spatial mode (first row, Figure 5) therefore marks ROIs where the variance-normalized signal changes little across the measured period. In the CSF, these are the infratentorial SAS and particularly the pontine cistern, where tracer enters from the spinal canal and enrichment is consistently high at most time points. In the parenchyma, deep white matter structures dominate for the opposite reason: these regions receive the least tracer within 24 hours.

#### 3.1.2 Component 2: The 24-hour supratentorial component

The 24-hour supratentorial component (component 2, second row Figure 4) has an increasing time profile dominated by the 24-hour peak, with no significant difference between patient groups. Its spatial mode (second row Figure 5) has high coefficients throughout the supratentorial SAS and in cerebral gray and white matter, with lower but still elevated values in the limbic system and basal ganglia, and near-zero values in the ventricles and infratentorial SAS. By 24 hours after injection, tracer will have spread from the infratentorial SAS into the supratentorial SAS and brain tissue in most patients, establishing a centripetal pattern (Ringstad et al., 2018; Solheim et al., 2026). Deep brain regions therefore participate in this distribution as well, despite receiving lower absolute enrichment than cortical gray matter. Because this component ties CSF and parenchymal ROIs to the same temporal profile and the same subject weights, enrichment in the two compartments varies together across subjects. Coupled enrichment between CSF in the SAS and ISF in the parenchyma is a central premise of the glymphatic framework (J. Iliff et al., 2012) and has been observed previously in humans (Ringstad et al., 2018; Wåhlin et al., 2026). This component provides a direct quantification of that relationship.

#### 3.1.3 Component 3: The early stage tracer influx component

The early-influx component (component 3, third row Figure 4) has a time profile that increases to a peak at 8 hours and returns to zero at 24 hours. The subject mode shows a statistically significant difference between the two groups (Mann–Whitney *U* test, *p <* 0.0001), with higher coefficient values in the REF group than in the iNPH group. The spatial mode (third row, Figure 5) has elevated coefficients in the infratentorial SAS, particularly the pontine cistern, and in the supratentorial SAS close to the frontal lobe. In the parenchyma, cortical gray matter in the frontal lobe (particularly the prefrontal cortex) and temporal lobes is prominent, together with the insula and subcortical structures in the medial temporal lobe, limbic system, and basal ganglia. We interpret this component as an influx pattern, characterizing tracer transport from infratentorial CSF into supratentorial regions in the SAS and brain tissue. The involved regions are consistent with transport along major cerebral arteries, which has been proposed as a preferential pathway for tracer transport in humans (Eide & Ringstad, 2024; Yamamoto et al., 2024). The transient time profile is also consistent with reports that this route contributes mainly to initial influx (Causemann et al., 2026; Eide & Ringstad, 2024).

#### 3.1.4 Component 4: The ventricular reflux component

The ventricular reflux component (component 4, fourth row Figure 4) shows a similar time profile to the early-influx component, with an increasing value up to 8 hours after injection and a subsequent decrease. Unlike the early-influx component, the time coefficient at 24 hours does not decay to zero, but rather remains at an elevated value. In the subject mode we find a statistically significant difference between the REF and iNPH groups (Mann–Whitney *U* test, *p <* 0.0001), with higher values in the iNPH group. The spatial mode (fourth row Figure 5) shows particularly elevated coefficients in ventricular CSF, as well as some more limited involvement of ROIs in the infratentorial SAS. In the parenchyma, ROIs in periventricular white matter, the basal ganglia and some parts of the medial temporal lobes are also involved. The consistently high coefficients in ventricular CSF indicate that this component reflects tracer entering ventricular CSF, a pattern known as ventricular reflux. The elevated coefficients in periventricular regions also suggest that tracer moves across the ventricular wall into the parenchyma. Retrograde flow with tracer entering ventricular CSF is a recognized feature of iNPH (Lindstrøm et al., 2018), and ventricular reflux has previously been graded in gMRI studies of iNPH patients (Eide et al., 2020).

### 3.2 REF and iNPH patients display distinct tracer influx routes

In the 4-component model, the early-influx and ventricular reflux components separate the REF and iNPH groups in opposite directions: early-influx is expressed more strongly in REF subjects, ventricular reflux in iNPH subjects (Mann–Whitney *U* test, *p <* 0.0001, Figure 4). Both have a time profile consistent with tracer influx, peaking at 8 hours before decreasing towards 24 hours, but the early-influx weight returns entirely to zero while the ventricular reflux weight remains at *∼* 0.5. The clearest difference between the two is in the spatial mode. Both have elevated coefficients in the infratentorial SAS, but ventricular CSF dominates the ventricular reflux component while being near-zero in the early-influx component. The parenchymal patterns also differ: ventricular reflux primarily involves periventricular regions, early-influx mainly cortical and some subcortical ROIs.

### 3.3 Ventricular reflux is correlated with age

In Figure 6 we investigate whether the subject mode of any of the components is significantly different based on diagnosis and if they are correlated with subject age, height or weight. We estimate correlations in the two diagnosis groups separately due to the group differences we observe in some of the components. The consistent observation from Figure 6 is that the CP components are largely independent of age, height and weight. The only significant correlation we find in this analysis is a positive correlation between age and the ventricular reflux component, which is present in both REF (*R*^2^ = 0.28*, p* = 0.0005) and iNPH patients (*R*^2^ = 0.33*, p* = 0.02), indicating that older patients may be slightly more likely to experience ventricular reflux.

**Figure 6:**
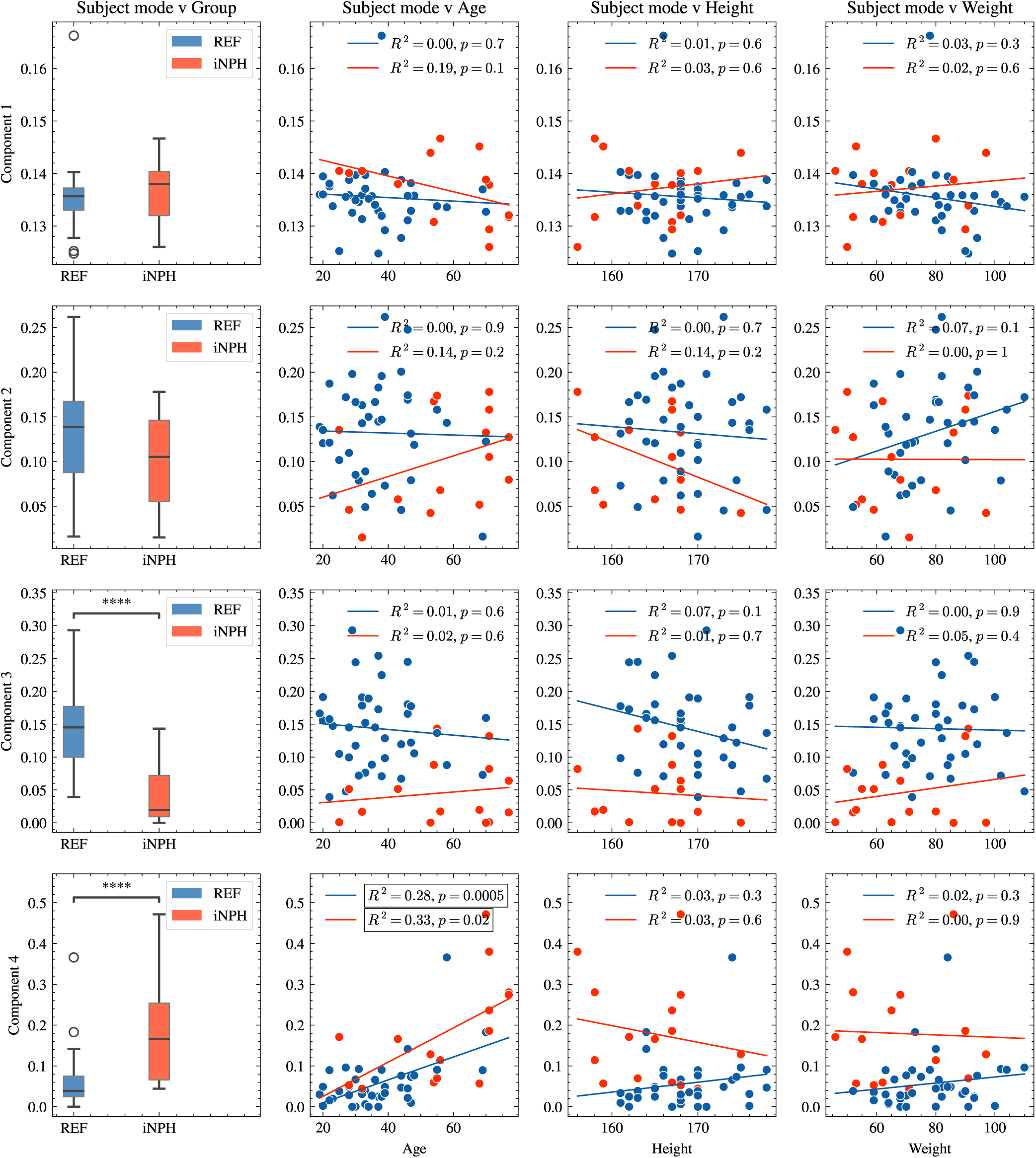
Subject mode coefficients obtained for each component as a function of various subject characteristics. From left to right: diagnostic cohort comparisons (iNPH vs. REF; Mann–Whitney *U* test, *∗ ∗ ∗∗ p <* 0.0001), age, height, and weight. Linear regressions were evaluated separately for iNPH (red) and REF (blue) groups (*R*^2^ and *p*-values computed via Wald test). Component 4 (ventricular reflux) demonstrates a statistically significant positive correlation with age across both cohorts (*p <* 0.05), whereas body height and weight show no significant relationships across any components.

### 3.4 Early-stage tracer influx is associated with later stage tracer enrichment

In Figure 7 we investigate the relationship between the components by computing the correlation between subject modes. We find that multiple of the components are correlated, indicating that several of the tracer patterns we identify may influence each other.

**Figure 7:**
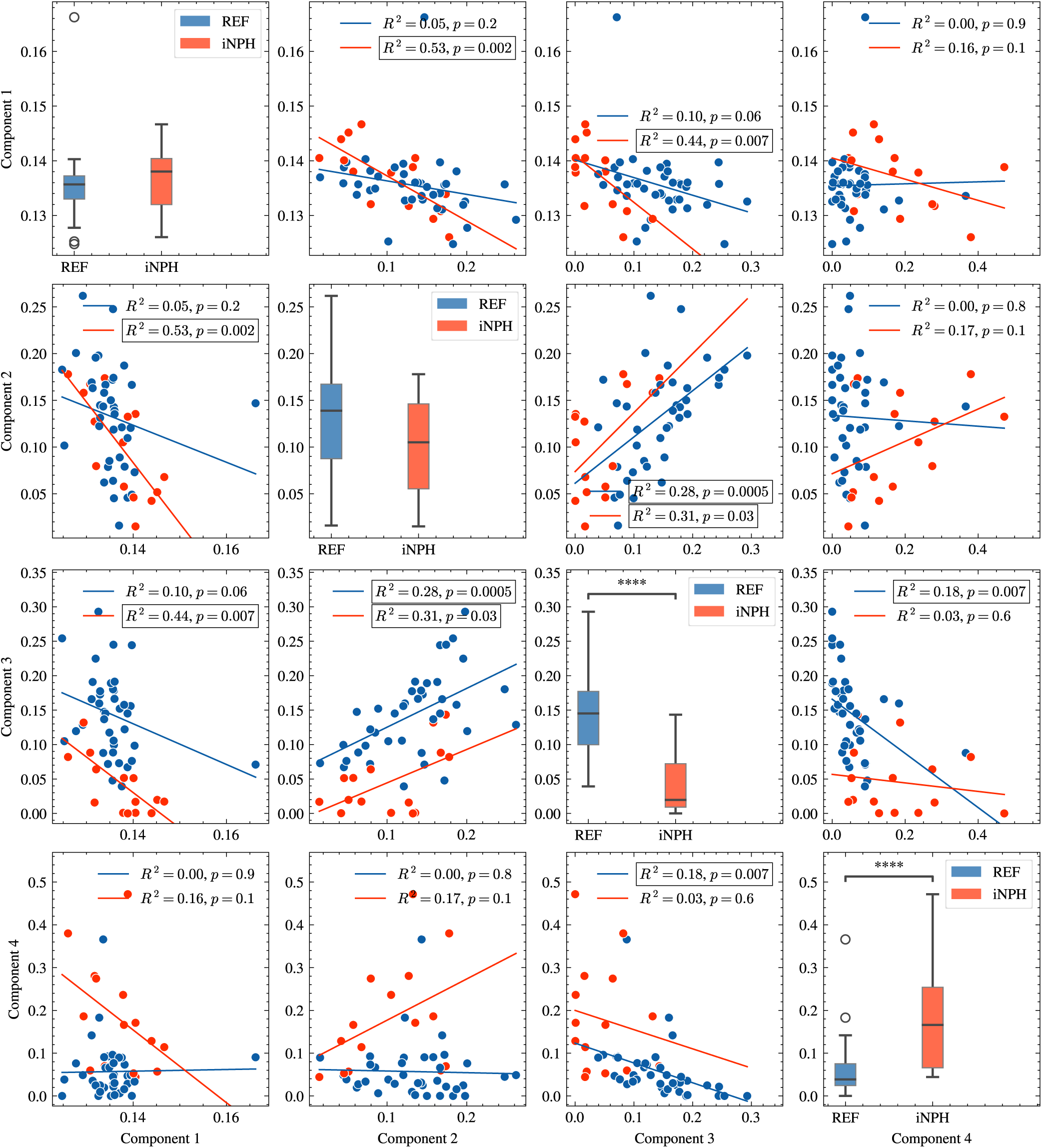
Pairwise correlations of subject mode coefficients across the four CP components in the female dataset; components 1–4 are the static, 24-hour supratentorial, early-influx, and ventricular reflux components respectively. Scatter plots and linear regression lines are shown separately for iNPH (red) and REF (blue) subjects. Statistically significant relationships (*p <* 0.05, Wald test) are highlighted with outlined legend boxes. Components 2 and 3 are positively correlated in both groups (*R*^2^ = 0.31*, p* = 0.03 in iNPH; *R*^2^ = 0.28*, p* = 0.0005 in REF). Component 1 is negatively correlated with components 2 (*R*^2^ = 0.53*, p* = 0.002) and 3 (*R*^2^ = 0.44*, p* = 0.007) in the iNPH group. In the REF group, components 3 and 4 are negatively correlated (*R*^2^ = 0.18*, p* = 0.007).

The most consistent correlation we find is between the 24-hour supratentorial component and the early-influx component, which is statistically significant in both REF and iNPH patients. The positive correlation between these two components suggests that the role of early-influx is to transport tracer from the infratentorial SAS, where tracer enters the brain, into the supratentorial SAS and cerebral brain tissue. We are accordingly able to directly quantify the relationship between early-stage dynamics and late-stage 24 hour dynamics captured by the 24-hour supratentorial component.

In addition, we find a statistically significant negative correlation between the static component and both the 24-hour supratentorial and early-stage components in the iNPH group, which is not present in the REF group. Finally, we find that early-stage and ventricular reflux components are negatively correlated in the REF group, indicating that the two routes may mutually compensate each other to some degree in this group.

### 3.5 CP components provide a quantitative grading of tracer transport

In a previous study using gMRI, patients were graded based on the degree of ventricular reflux (Eide et al., 2020), which is a grading scale that targets a pattern similar to the ventricular reflux component we derive in this work. The main difference is that while grading in (Eide et al., 2020) is discrete and based on expert raters, the CP component provides a data-driven and continuous grading scale. Figure 8 shows an example of how patients can be graded on this scale, using the associated subject coefficient *c*_4_. In this figure we show the time evolution of three subjects: the first subject has a median ventricular reflux score for iNPH patients (*c*_4_ *≈* 0.2); the second subject has a median score for REF patients (*c*_4_ *≈* 0.05); the third subject is an extreme case in the REF group with a score of *c*_4_ *≈* 0.2. We note that the median iNPH patient, in the left column, has substantial tracer enrichment in the ventricle while the median REF patient has no signal increase in this region. Meanwhile, the REF outlier clearly shows considerable signal increase in the ventricle. This patient therefore receives a score comparable to the median iNPH patient, showing that *c*_4_ varies continuously across the cohort rather than separating along diagnosis.

**Figure 8:**
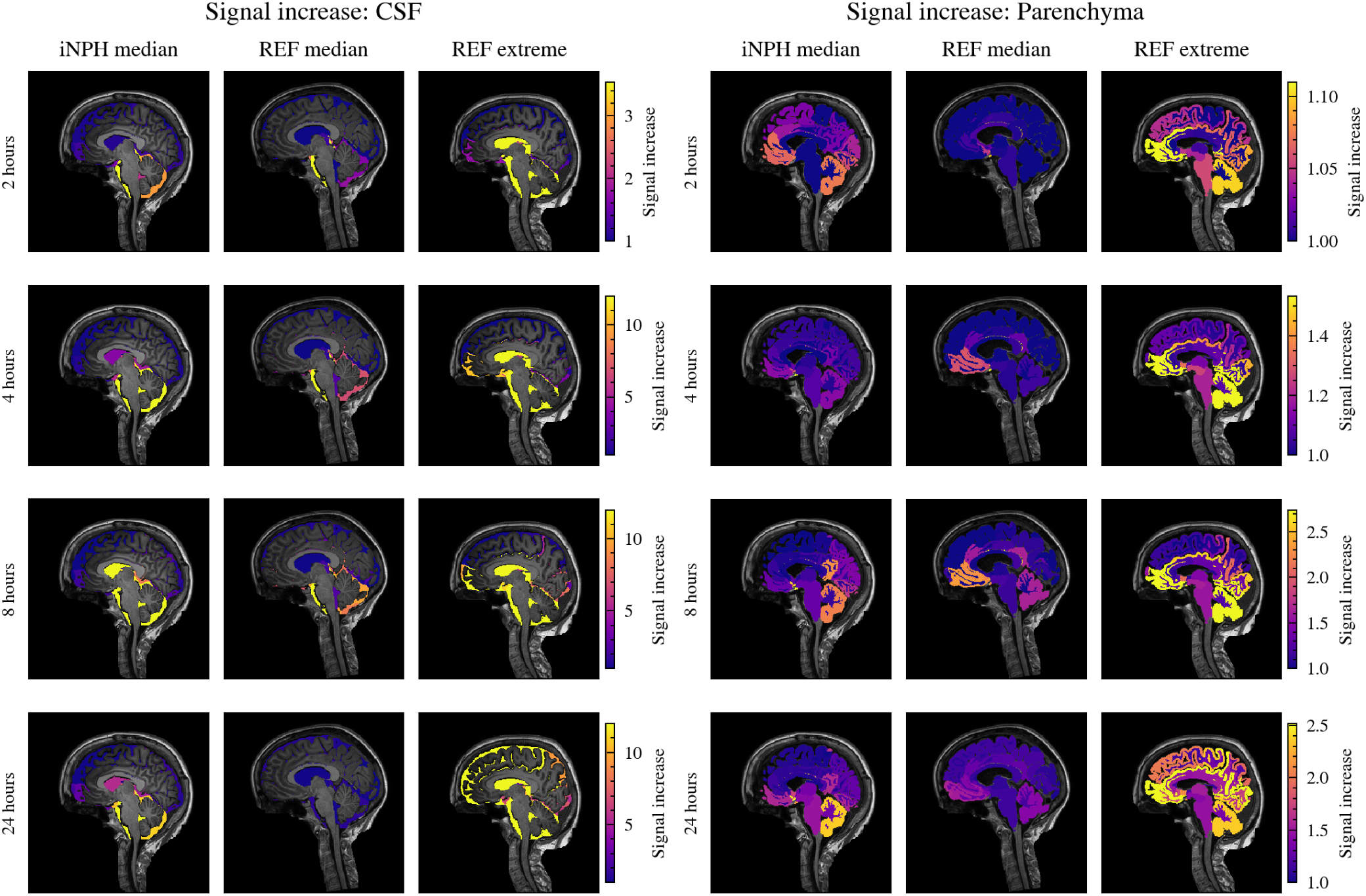
Longitudinal *T*_1_ signal increase across CSF and parenchymal compartments in three representative subjects, illustrating the ventricular reflux grading provided by the ventricular reflux component subject coefficients *c*_4_. Left: A typical iNPH patient with a median score in that cohort (*c*_4_ *≈* 0.20), exhibiting prominent, sustained ventricular reflux. Middle: A typical REF control subject with a median score (*c*_4_ *≈* 0.05), showing no tracer entry into ventricular CSF. Right: An outlier in the REF group (*c*_4_ *≈* 0.20), demonstrating marked ventricular tracer enrichment and elevated parenchymal signal comparable to the iNPH cohort.

## 4 Discussion

Treating tracer signal in CSF spaces and brain parenchyma within a single model allows coupled spatiotemporal patterns to be recovered without *a priori* spatial assumptions. From the female subset we obtain four replicable and interpretable components, three of which correspond to identifiable transport features, and whose subject-specific weights can be compared directly against one another within the same individuals. The fourth, the static component, is not interpreted as a transport pathway, but rather as a result of modeling choices discussed in Section 4.6. We discuss each component and the relationships between them below, before considering what such components can and cannot establish about the underlying physiology.

### 4.1 Quantifying CSF–ISF coupling and tracer influx

The coupling between CSF and ISF is a central claim of the glymphatic theory (J. Iliff et al., 2012), which proposes that CSF enters brain tissue through periarterial spaces and exits along perivenous spaces. While the mechanics of solute transport in the brain remain debated (Kipnis et al., 2025; Mestre et al., 2020), the direct coupling between CSF in the SAS and ISF in the parenchyma is central to how solutes are cleared from the brain.

Several of the components we compute in this work are consistent with the notion of a direct coupling between tracer enrichment in the SAS and regions deep within brain tissue, with the 24-hour supratentorial component being the clearest example. This component shows areas of the supratentorial SAS that evolve with the same time profile as regions of cerebral gray and white matter, thereby directly coupling CSF in the SAS to ISF in the parenchyma. As previously mentioned in Section 3.1.2, this coupling has also been noted in previous studies of gMRI in humans (Eide & Ringstad, 2015; Ringstad et al., 2018; Wåhlin et al., 2026), but it has not been easily quantifiable in the way enabled by the present approach. Because each component carries a subject-specific weight, its expression can be compared directly against the other transport patterns in the same individuals. An additional key observation from Figure 5 is that the coefficient values are elevated within subcortical white matter, not only gray matter, indicating that these regions are also involved in CSF–ISF interaction. While these are regions that generally receive little tracer enrichment in absolute terms, their relative involvement appears substantial 24 hours after injection.

The early-influx and ventricular reflux components also show the direct CSF–ISF interaction involved in tracer transport. In the early-influx component we note the elevated coefficients in several cortical gray matter regions and several subcortical structures, particularly in the medial temporal lobes, closely reflecting the patterns we observe in the SAS. The ventricular reflux component also shows that periventricular regions have elevated coefficient values, indicating that tracer appears to cross the ventricular ependyma into the parenchyma. This component therefore ties the ventricular CSF and the surrounding tissue to the same temporal profile. Both of these components accordingly show that tracer in the SAS and in the parenchyma evolve in concert, underpinning the role of CSF–ISF interaction.

While the early-influx component involves CSF–ISF interaction, its main feature is another key part of how tracer distributes itself in the brain: transport along major cerebral arteries. Recent work using intrathecal contrast agents has suggested that perivascular spaces, surrounding major cerebral arteries, act as a preferential pathway for tracer transport in humans (Eide & Ringstad, 2024; Yamamoto et al., 2024). The pattern captured by the early-influx component is consistent with this pathway, showing elevated coefficients in regions that correspond to the locations of the anterior and middle cerebral arteries. The time profile suggests that this is a transient phenomenon, with an effectively zero weight 24 hours after injection, which matches observations that this is primarily a mechanism for initial tracer influx (Causemann et al., 2026; Eide & Ringstad, 2024). We emphasize, however, that at the ROI resolution used here perivascular spaces cannot be resolved, meaning that the component is consistent with transport along major arteries rather than evidence for it.

### 4.2 REF and iNPH patients display distinct tracer influx routes

The clearest distinction between REF and iNPH patients in this study lies in the route of early tracer influx. Both the early-influx and ventricular reflux components describe tracer entering the SAS and brain tissue from the spinal canal, and both are expressed in both cohorts, but ventricular reflux is more strongly expressed in iNPH and early-influx in REF. Ventricular reflux is a hallmark of the iNPH diagnosis: these patients suffer from CSF flow disruptions that can produce retrograde flow in the cerebral aqueduct (Lindstrøm et al., 2018), causing tracer to enter ventricular CSF. The contribution of the present analysis is that the alternative route is quantified alongside it: the early-influx component describes transport from the infratentorial into the supratentorial SAS in regions consistent with transport along major cerebral arteries. The separation is not a clean classification, however. The subject mode distributions overlap considerably, particularly for ventricular reflux, which may reflect measurement resolution, segmentation errors, or the substantial individual variation in tracer arrival time reported in gMRI studies (Eide, Mariussen, et al., 2021; Hovd et al., 2022). The REF and iNPH diagnoses are also not based on CSF flow dynamics alone and therefore an unsupervised decomposition would not be expected to reproduce them exactly.

Although we find that the early-influx pattern has a lower expression among iNPH patients, it does not preclude the presence of transport along major arteries in this group. Previous studies using gMRI have shown that iNPH patients still display tracer transport along this route, but more slowly than in the REF group (Eide & Ringstad, 2024). The ventricular reflux spatial mode also does not preclude transport along major arteries: it has elevated coefficients in the medial temporal lobe and in parts of the temporal cortex and insula, regions that are also prominent in the early-influx component. The group difference we observe may therefore be due to CP being too restrictive by making all subjects share the same time pattern. Patients with iNPH may express a pathway similar to early-influx with a delayed temporal pattern that a single set of time coefficients cannot represent. Decompositions such as PARAFAC2 (Mørup et al., 2025), which allow subject-specific time profiles, would be needed to distinguish these. It is also worthwhile to note that, in the REF group, the early-influx and ventricular reflux components are negatively correlated, indicating that strong expression of one route accompanies weaker expression of the other. We note, however, that this relationship accounts for a small fraction of the variance and is not observed in the iNPH group, where the smaller subset size limits what can be inferred.

Finally, we note that the time modes for the early-influx and ventricular reflux components display a similar influx pattern peaking at a value in the range 0.7 *−* 0.8 at 8 hours and then decreasing towards 24 hours. The main difference is that while the early-influx time mode goes essentially to zero, the ventricular reflux component remains substantial. This is consistent with delayed tracer clearance in patients with a strong expression of this component, which matches gMRI studies reporting delayed brain clearance in iNPH patients (Eide & Ringstad, 2018; Ringstad et al., 2018). The sustained coefficient at 24 hours likely has at least two contributions. The primary one is slow clearance from ventricular CSF itself: if retrograde flow allows tracer to enter the ventricles, the same disrupted dynamics could plausibly also delay tracer from exiting that same region. A secondary contribution is tracer that crosses the ventricular ependyma into periventricular white matter, where transport is slower than in CSF and clearance correspondingly delayed. The tracer enrichment in tissue will however be much lower than in CSF. The prominence of periventricular ROIs in the spatial mode reflects their relative involvement after variance normalization rather than comparable absolute tracer concentrations.

### 4.3 Early-stage tracer influx is associated with later stage tracer enrichment

The components allow us to directly couple early stage tracer influx with later stage enrichment in a quantitative manner. Figure 7 shows that the 24-hour supratentorial component and the early-influx component are positively correlated in both the REF and the iNPH groups. This supports the view of the early-influx component as an influx pattern, responsible for transport of tracer from the infratentorial SAS into supratentorial regions, which directly enables the tracer to reach the regions involved in CSF–ISF coupling. The early-influx component differs between diagnosis groups while the 24-hour supratentorial component does not, despite the correlation between them. Although the REF median is higher, the spread is still too wide for the difference to reach significance. This may reflect individual variation in tracer arrival time, since at 24 hours some patients may already be in the decline phase in superior regions while others are still enriching. Additional later time points would help separate variation in arrival time from lower total enrichment.

Additionally, we find that in the iNPH group the static component is negatively correlated with the 24-hour supratentorial and the early-influx component. This negative correlation may be partly due to components not being orthogonal: when the time modes show conflicting patterns (static versus not static) we may expect the components to be anti-correlated when some of the regions with elevated coefficients overlap, as is the case for these three components (see Figure 5). The lack of a statistically significant relationship between the static component and the 24-hour supratentorial and early-influx components in the REF group appears to be driven by a single REF outlier in the static component. Removing this patient leads to a statistically significant correlation also in the REF group.

### 4.4 Ventricular reflux is correlated with age

The results in Figure 6 do not show any correlation between the four components and age, height, or weight, except in the ventricular reflux component. In this case, we find a positive correlation between age and the subject coefficient which is statistically significant for both REF and iNPH patients. Because iNPH patients in this dataset are consistently older than REF subjects, the age effect could in principle reflect the demographic imbalance. The fact that the correlation exists within each patient cohort separately suggests that this is indeed an age related effect. The trend of older patients being more likely to experience ventricular reflux may in part be associated with brain atrophy, which leads to increasing ventricular volume in humans with age (Fjell et al., 2013; Jack et al., 2008; Madsen et al., 2015). Large ventricular volumes may alter CSF dynamics, potentially allowing tracer to enter ventricular CSF, even in non-pathological cases. However, from this present dataset it is hard to completely disentangle age-related brain atrophy from ventriculomegaly due to CSF flow disorders associated with iNPH. Studies grading the amount of ventricular reflux in the REF and iNPH cohorts also find a higher prevalence of this pattern in the iNPH group (Eide et al., 2020).

The lack of a statistically significant relationship between any of the components and factors such as height or weight could mean that body size has a minimal effect on how the tracer itself spreads within the brain. However, in this study we are not directly controlling for the absolute amount of tracer that enters the brain, which would require quantitative MRI (*T*_1_ maps). It is still possible that the amount of tracer that arrives in the brain following intrathecal injection could vary substantially based on parameters like height and weight.

### 4.5 CP components provide a quantitative grading of tracer transport

Subject coefficients of a CP component offer two distinct advantages over grading scales from expert raters. First, they provide a continuous scale rather than a small number of ordinal grades. As an example, the grading in (Eide et al., 2020) involves five discrete steps; by using a continuous scale we can classify subjects with greater granularity. Second, the score reflects each subject’s expression of the full spatiotemporal pattern, including periventricular parenchyma and the temporal profile out to 24 hours, rather than the appearance of a single region at a single time point. The same criterion is applied to every subject without rater judgment, which may provide a more consistent basis for classification.

The caveat of using CP for this purpose is that we do not know exactly what the component is identifying. We ascribe an interpretation to the pattern we observe, but the component may also include features which are less relevant for clinical purposes. If the analysis is restricted to only considering the extent of tracer enrichment in the ventricles, then an approach targeting those few ROIs specifically may be preferable. However, in many cases the global patterns which give rise to characteristic properties of a given patient group can be hard to determine *a priori*. In this case unsupervised approaches may provide a data-driven alternative to determine composite regions of interest. Finally, we note that the subject coefficients are estimated jointly with the spatial and time modes on this cohort. Applying the grading to a new patient would require projecting that subject onto the fixed modes, which we have not tested here.

### 4.6 What CP components can and cannot establish

While CP components are derived in an unsupervised manner without *a priori* knowledge of the patterns we expect, the outcome of the decomposition is still subject to distinct modeling and data processing choices. Interpretations are made *post hoc* and are not necessarily tied to any single physiological mechanism because they are the result of a fitting process. The clearest example of this is the static component, which corresponds essentially to an average. It would be possible to remove this average component from the decomposition by centering the data prior to computing the CP, i.e., subtracting the average of all time points and subjects from each spatial slice. In this work, we have chosen not to do this in order to maintain non-negativity. Because tracer enrichment is estimated by dividing the post-injection MRI by the pre-injection MRI, the data itself is inherently non-negative and components are therefore easier to interpret when we retain this property. Additionally, non-negativity can act as an effective regularization of the objective function which can be beneficial for the uniqueness properties of the solution. This component should therefore not be interpreted as a tracer pathway, but as a consequence of the non-negativity constraint together with the variance scaling we use to ensure more consistent weighting of ROIs.

The design of the CP model itself may also be too restrictive to capture all relevant patterns in the data. A single set of time coefficients is shared by all subjects, yet previous studies report substantial variation in the time at which tracer concentration peaks (Eide, Mariussen, et al., 2021; Hovd et al., 2022). Decompositions such as PARAFAC2 (Mørup et al., 2025), which allow subject-specific time profiles, would account for this directly, and might also better isolate the spatial distribution of the tracer and differences between groups while reducing the influence of variation in arrival time.

Another key restriction of the components we derive is the need for a shared spatial representation. In this work, we use patient specific ROIs generated using FreeSurfer (Dale et al., 1999; Desikan et al., 2006), and then take the median signal in each region to generate the tensor. While each image, at each time point, will have on the order of 10^6^ voxels with relevant signal this averaging reduces the spatial resolution to 245 ROIs. In combination with the relatively low temporal resolution of the longitudinal MRIs, this means that there are many fine-grained details we cannot resolve in this analysis. We can for instance not separate the role of peri-arterial versus peri-venous transport or the effect of higher frequency pulsations in tissue or CSF due to breathing or vascular effects.

A common challenge in longitudinal studies of tracer clearance in humans and animals is how to robustly interpret the data observed clinically and experimentally in order to understand efflux pathways. In studies involving tracers injected intrathecally or in the cisterna magna (in rodents), tracer influx to the parenchyma will be happening simultaneously as efflux back into CSF (Wåhlin et al., 2026) and from CSF into dural lymphatics. Unlike rodents, which generally display efflux to olfactory regions (Kroesbergen et al., 2024; Sigurdsson et al., 2023), humans do not appear to have one single solute efflux pathway from CSF into dural lymphatics or from parenchyma into CSF. The combined influx and efflux process makes individual clearance pathways hard to identify from the data itself, a challenge that CP does not overcome. For instance, the static component shows that pontine cistern enrichment varies little across the measured period, meaning tracer remains in this region at 24 hours, well after the initial infratentorial influx phase. One explanation is continuous local efflux, which would be consistent with the recent evidence for clearance along cranial nerves (Falkenberg-Jensen et al., 2025). However, sustained signal is equally compatible with continued arrival of tracer from the spinal canal, and the component alone cannot distinguish the two. Using higher resolution spatial maps and including later time points, when the tracer will in most cases have fully cleared from the SAS, may make this possible to resolve.

### 4.7 Limitations and further work

The dataset analyzed in this work is large for a longitudinal study of this kind. It is rare to have access to longitudinal MRI datasets with close to 100 patients. However, given the well-known individual variations in gMRI studies (Eide, Mariussen, et al., 2021; Hovd et al., 2022) and the relative imbalance between sex, diagnosis, and age among patients in the dataset, the number of patients is a limiting factor in drawing definitive conclusions from this work. Additional data with improved balance within key characteristic groups may be needed to derive components which can effectively be translated into a clinical setting. Still, we emphasize that tensor-based methods may be a critical tool in combining data from different modalities recorded on the same patient population (Mørup et al., 2025). Such tools could also leverage large low-information datasets to extract information from smaller high-detail datasets, such as the one in this work, through foundation models (Dong et al., 2025; Jiang et al., 2026).

Future work should also aim to make better use of all the available spatial information. In this present work, we use segmentations that result in relatively coarse spatial averaging. An alternative approach would be to map all patients to a common geometry using image registration. Previous studies have shown how this can be done on gMRI data (Solheim et al., 2025, 2026). However, doing this would also significantly increase the computational effort of computing the decomposition.

Finally, errors due to partial volume effects and segmentation errors can never be fully excluded when working with longitudinal tracer data. Patients with enlarged ventricles are often particularly difficult to accurately segment using automated segmentation software. Particularly ROIs close to the ventricles may be influenced by this effect. In addition, partial volume effects can significantly alter the signal in a given voxel at the interface between tissue and CSF. The tracer signal increase in the CSF will often be much larger than in the parenchyma, meaning that errors in the segmentation at this interface may importantly alter the estimated signal. In this work, we aim to limit this effect by taking the median signal increase in any given ROI in order to reduce the impact of outlier signal increases. However, it is possible that some of the increased coefficients we observe in the ventricular reflux component close to the ventricles could be particularly exposed to signal leakage.

## 5 Conclusion

This study demonstrates the efficacy of tensor low-rank approximations, specifically the Canonical Polyadic (CP) decomposition, for analyzing complex, multi-subject longitudinal imaging data of solute transport in the human brain. We perform a stratified analysis of the dataset focusing on the female subset, from which we extract four robust and replicable components that capture distinct tracer fluid pathways. By simultaneously treating tracer signal in the CSF and parenchyma we identify a component that shows the direct coupling between CSF in the SAS and ISF in the parenchyma. Another component captures the early stage dynamics of tracer, illustrating the involvement of regions consistent with transport along major cerebral arteries. This component is found to be positively correlated with 24 hour distribution, establishing a direct relationship between tracer influx and later stage enrichment. We also find a component showing ventricular reflux, where tracer enters ventricular CSF. Its expression is significantly elevated in iNPH relative to REF subjects, it correlates positively with age, and it provides a continuous, rater-independent grading of a pattern currently scored by expert consensus. While current limitations in spatial resolution restrict our ability to definitively pinpoint specific efflux pathways, this work offers an approach for directly quantifying complex interactions in solute transport. As research into the glymphatic system expands, leveraging tensor methods could be the key to managing high-dimensional, multimodal neuroimaging data and developing robust, image-based biomarkers for clinical diagnostics.

## Declarations

### Funding

K.A.M. and A.S. acknowledge funding by the European Research Council under grant 101141807 (aClean-Brain) and the national infrastructure for computational science in Norway, Sigma2, via grant NN9279K. M.E.R has received funding from the Research Council of Norway (RCN) via grant #360005 (DigiCells). This work was supported by the foundation Stiftelsen Kristian Gerhard Jebsen through its program for translational medical research via the K. G. Jebsen Centre for Brain Fluid Research, and by the Center of Advanced Study at the Norwegian Academy of Science and Letters under the program Mathematical Challenges in Brain Mechanics.

## Acknowledgements

Computational experiments and data processing were performed on resources provided by Sigma2 —the National Infrastructure for High-Performance Computing and Data Storage in Norway, under grant NN9279K. A.S. thanks Lars M. Valnes for assistance in data preparation.

## Ethics and approvals

Parts of the data presented in this work have also been used in previous works on MRI-based assessment of human glymphatic function conducted at the University Hospital of Oslo (Ringstad et al., 2017, 2018) in the years 2015*−*2019. Collection of data analyzed for this study was approved by the Regional Committee for Medical and Health Research Ethics (REK) of Health Region South-East, Norway (2015/96), the Institutional Review Board of Oslo University Hospital (2015/1868) and the National Medicines Agency (15/04922-7), and was conducted following the ethical standards of the Declaration of Helsinki of 1975 (revised in 1983). Study participants were included after written and oral informed consent. No new data were collected for the present work.

## Competing interests

The authors declare no competing interests.

## Code and data availability

The dataset used in this paper is not publicly available due to patient data privacy concerns. The code for this work is made available at https://github.com/Erasdna/Tensor4gMRI.git. The time and spatial modes of the CP models presented in this work can be downloaded from https://doi.org/10.5281/zenodo.22669838. The subject mode is withheld due to patient privacy concerns.

## Declaration of the use of AI

The AI models Gemini Flash 3.6 and Claude Sonnet 5 were used for editing and revision of this manuscript to identify language and spelling errors and to improve clarity. All AI generated text was checked and verified by humans.

## CRediT

**Andreas Solheim**: Writing: Original draft, Review & editing. Conceptualization, Methodology, Software, Investigation, Visualization. **Geir Ringstad**: Writing: Review & editing, Data collection and curation. **Per Kristian Eide**: Writing: Review & editing, Data collection and curation. **Marie E. Rognes**: Writing: Review & editing, Conceptualization. **Evrim Acar**: Writing: Review & editing, Conceptualization, Methodology. **Kent-Andre Mardal**: Writing: Review & editing, Conceptualization, Methodology, Supervision.

## Supplementary material: Stratified CP models of the dataset

To this point we have limited the analysis to the female subset of the dataset. Due to the correlation between sex and diagnosis among patients in the dataset, as seen in Table 1, we argue that this is the most reliable subset of the available subject population because this is the largest single group. However, splitting the dataset to include only males, REF or iNPH patients also leads to replicable results, as seen in Table 2. In this section, we provide the equivalent figures to Figures 4, 5 and 6 for the largest number of replicable components we compute. Similar to the female subset, we compute the relative importance *λ_r_* of each component and then sort the components by decreasing value of *λ_r_*. We briefly summarize the findings from each of the models from the different subsets:

### 3-component model using Male patients

In the male subset we obtain three replicable components, largely consistent with the static, 24-hour supratentorial and early-influx components in the female subset. Component 1 in Figure S1 has a constant time profile and no significant difference between the iNPH and REF groups, matching the static component. Its spatial mode (Figure S2) shares the elevated coefficients in the pontine cistern and basal ganglia, but the periventricular white matter regions prominent in the female subset are instead replaced by subcortical white matter. Component 2 corresponds to the 24-hour supratentorial component in the time and subject modes, with a similar spatial pattern to Figure 5 apart from lower coefficients in the posterior cortex. Component 3 corresponds most closely to the early-influx component, but also carries features of the ventricular reflux component. In particular, the spatial mode has elevated coefficients in ventricular CSF, and the time coefficient does not return to zero after 24 hours. Unlike the female subset, its subject mode shows no significant difference between REF and iNPH patients. None of the components differ significantly between diagnosis groups or correlate with age, height or weight, except for a negative correlation between component 1 and height among iNPH patients (Figure S3).

### 4-component model using iNPH patients

In the iNPH subset we find four replicable components, all with direct equivalents in the female subset and in the same order. Component 1 in Figure S4 has a constant time profile, matching the static component. Its spatial mode (Figure S5) differs from the female subset in the same way as component 1 in the male subset: the basal ganglia and subcortical white matter have elevated coefficients, while the periventricular white matter regions emphasized in Figure 5 do not. Components 2–4 do not importantly differ from their counterparts in Figure 4. Comparing male and female subjects within this cohort, none of the components show a statistically significant difference in the subject mode, and we find few significant correlations with age, height or weight (Figure S6). Components 3 and 4 correlate with age among male subjects.

### 4-component model using REF patients

Similarly to the iNPH split, the 4 components we identify in the REF split are equivalent to the components from the female subset. Components 1 and 2 in Figure S7 are essentially equivalent to the static and 24-hour supratentorial components in Figure 4. Meanwhile, components 3 and 4 appear in the opposite order in the REF and female subsets. The main observation from Figure S7 is, however, the statistically significant group differences between male and female patients in the subject mode of components 2, 3 and 4. In particular, we find that components 2 and 3 have a slightly higher expression in the female subset, while component 4 has a higher expression in the male subset.

### Summary and discussion

When stratifying the cohort into other subsets (Male, REF, iNPH), we largely get similar components as the female only split. The CP decomposition is invariant to permutation and therefore the specific numbering of the components does not bear any significance. Except for the male split, the CP decomposition finds one component that is constant in time, one component which dominates at 24 hours after injection and two influx components, where one is consistent with early-stage periarterial influx and the other reflects ventricular reflux. The ordering of the components based on the relative importance *λ_r_* also gives an ordering that is consistent with the female subset. In the male subset, the third component appears to be a combination of the early-influx and ventricular reflux components we find in the female subset. Because these two patterns are expressed more strongly in opposite cohorts in the female model, a merged component inherits both group effects, which may explain the absence of a significant REF–iNPH difference in the male subject mode. We note that separating the two routes therefore depends on the model order, although the split is recovered consistently in the three subsets where a 4-component model is replicable.

While the time modes are generally consistent across splits, some spatial modes differ in ways worth noting. In particular, the periventricular white matter regions with elevated coefficients in the static component of the female subset are replaced by subcortical white matter in component 1 of the male and iNPH subsets. Since the constant time profile marks ROIs where tracer concentration varies little over the measured period, periventricular regions appear in this component in the female subset largely because they receive little tracer within 24 hours. In cohorts where tracer does reach these regions, their signal varies over time and they are instead captured by the ventricular reflux component. Both the iNPH and male groups are plausibly more prone to ventricular reflux: it is a hallmark of iNPH, and the male subjects in this dataset are consistently older than the female subjects (median age 65 versus 38, see Table 1), which is associated with increased expression of ventricular reflux in Figure 6.

**Figure S1:**
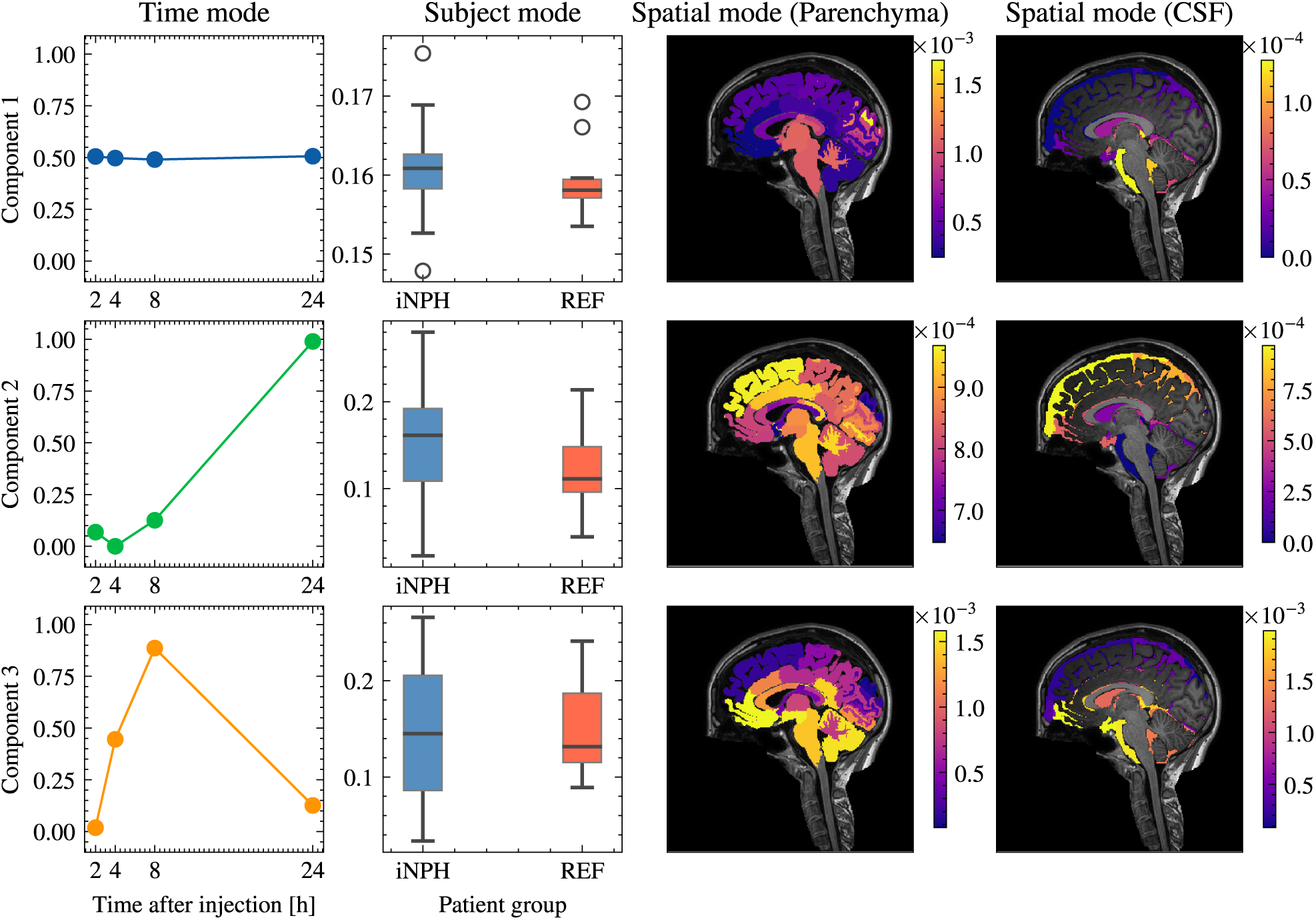
Time, subject and spatial modes obtained using a 3-component CP model on the male subset of the dataset. We split the spatial component into parenchyma and CSF to view the dynamics in each region more clearly.

The components we obtain on the REF subset are also worth specific scrutiny. Similarly to the female subset, we find a statistically significant difference between the two groups (male and female) in components 3 and 4. However, we note that the *p*-values are higher in the REF subset than in the female subset. While these results appear to indicate sex-specific differences in tracer transport, the dataset in this work is likely insufficient to conclude that this is the case. In particular, there is a significant age difference between the male and female patients in the dataset. Older patients experience brain atrophy, which leads to ventricular volume in humans increasing with age (Fjell et al., 2013; Jack et al., 2008; Madsen et al., 2015). Increasing ventricular volume could alter CSF flow, potentially making these patients more similar to iNPH patients in certain cases. An age- and sex-balanced dataset is necessary to establish whether there are any significant sex-based tracer transport differences.

**Figure S2:**
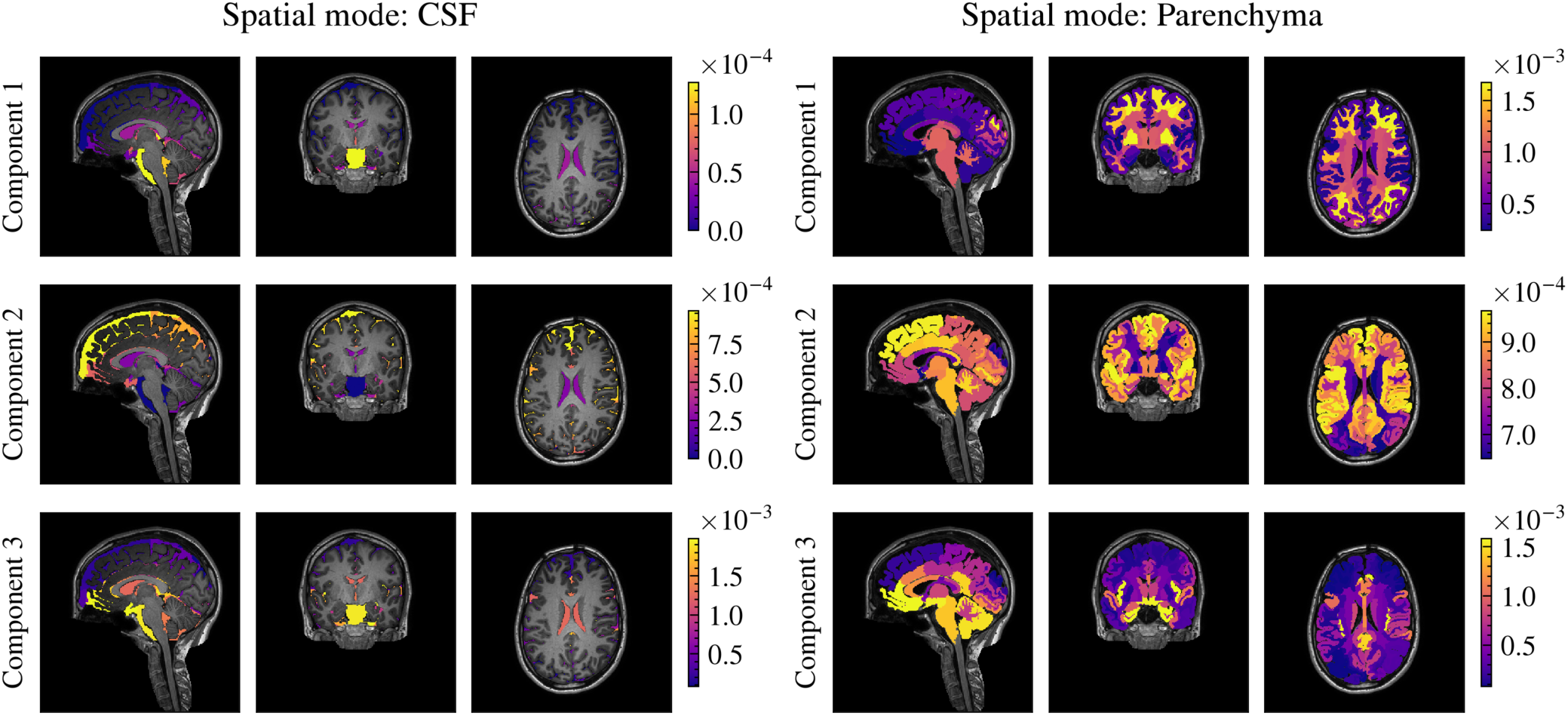
Sagittal, coronal and axial slices of the spatial mode for each component on the male subset of the dataset. Provides a detailed spatial view of Figure S1.

**Figure S3:**
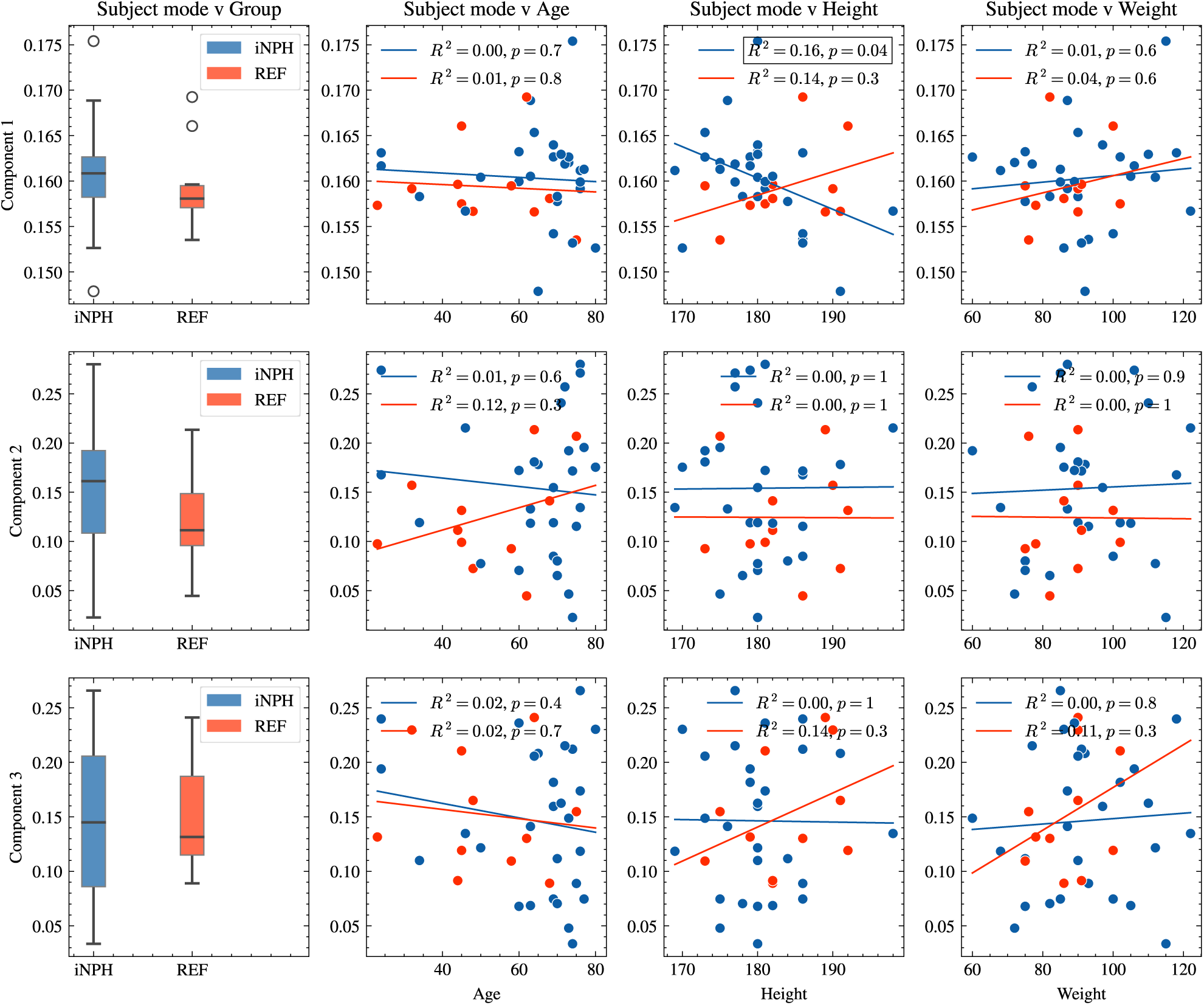
Subject mode coefficients obtained for each component as a function of various subject characteristics on the male subset of the dataset. We compare the two diagnosis groups (REF and iNPH), and find no significant differences for any of the components (Mann-Whitney U-test, *p <* 0.05). The subject mode coefficient is also plotted with respect to age, height and weight for each of the two groups.

**Figure S4:**
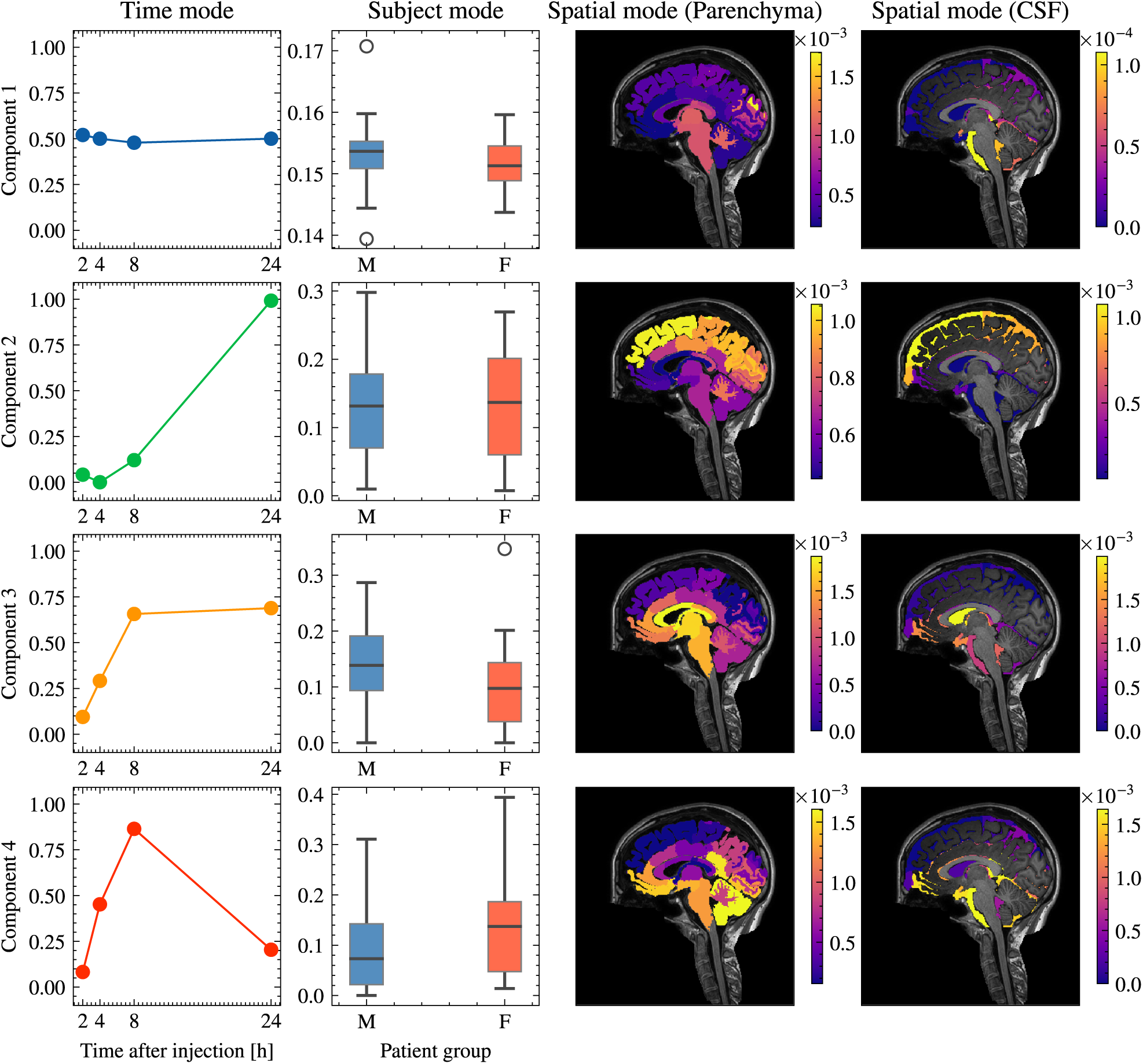
Time, subject and spatial modes obtained using a 4-component CP model on the iNPH subset of the dataset. We split the spatial component into parenchyma and CSF to view the dynamics in each region more clearly.

**Figure S5:**
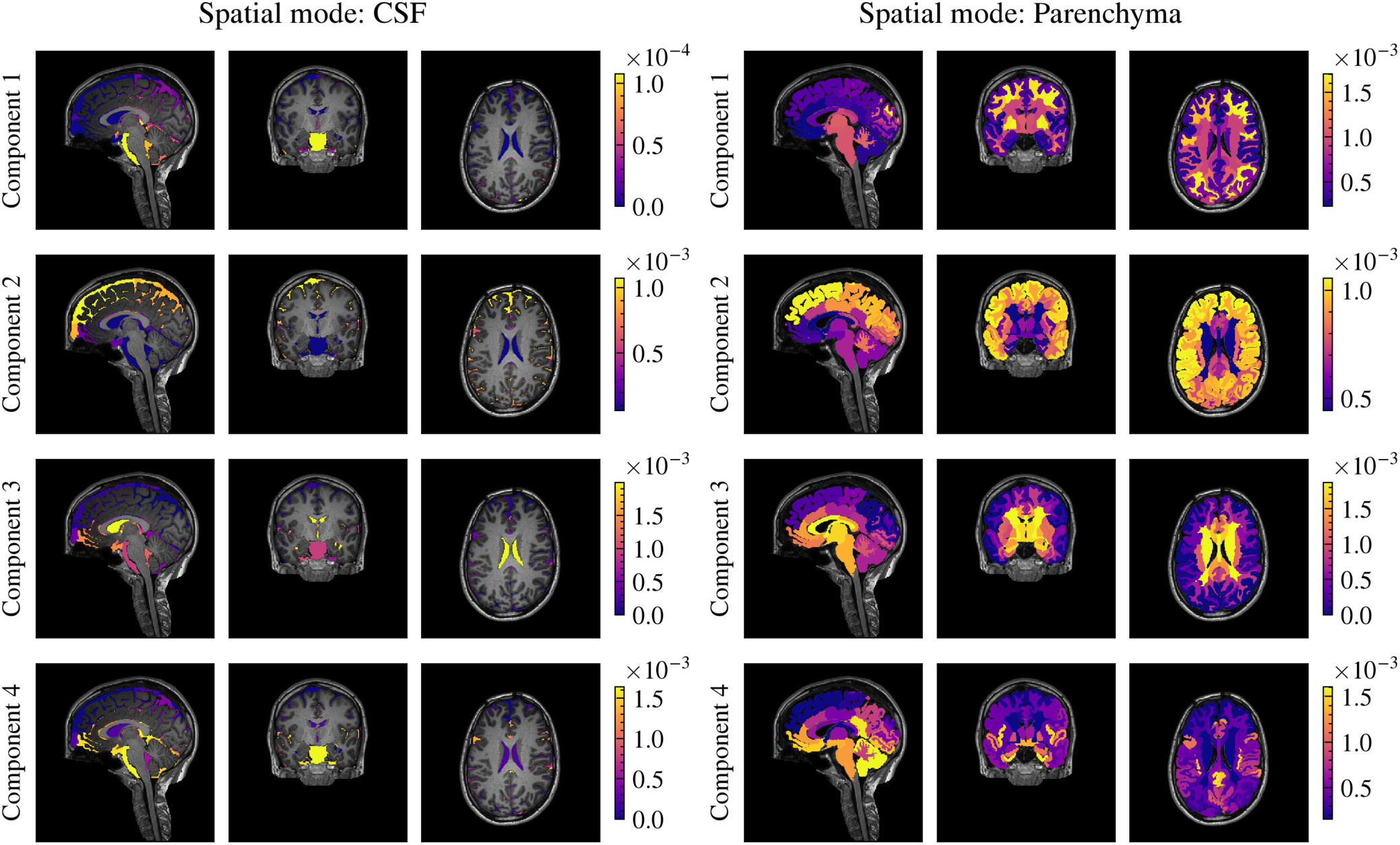
Sagittal, coronal and axial slices of the spatial mode for each component on the iNPH subset of the dataset. Provides a detailed spatial view of Figure S4.

**Figure S6:**
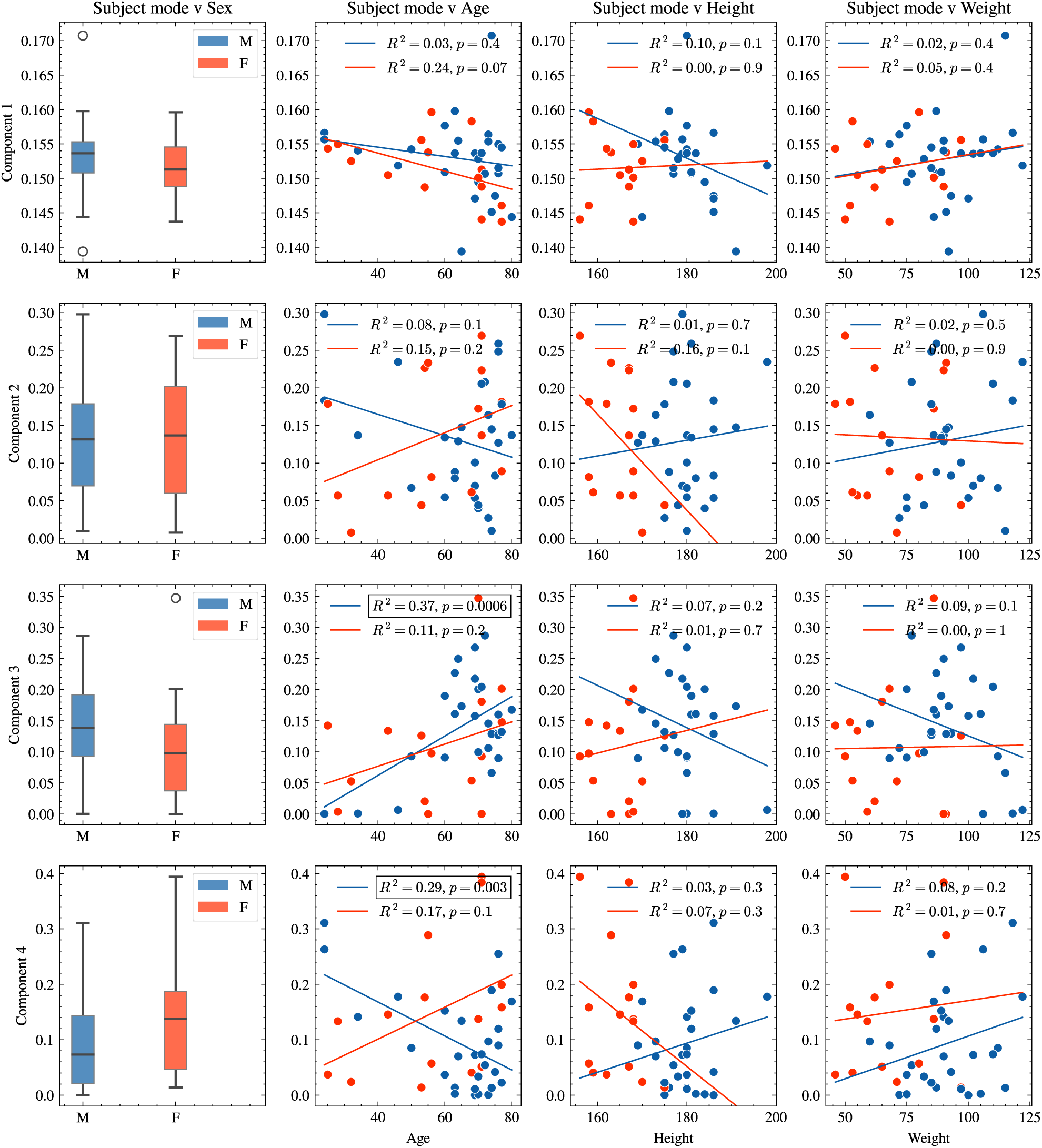
Subject mode coefficients obtained for each component as a function of various subject characteristics on the iNPH subset of the dataset. We compare the two groups (male and female), and find no significant differences for any of the components (Mann-Whitney U-test, *p <* 0.05). The subject mode coefficient is also plotted with respect to age, height and weight for each of the two groups. We find a significant relationship in the third and fourth components with regards to age among male subjects (Wald Test with t-distribution *p <* 0.05).

**Figure S7:**
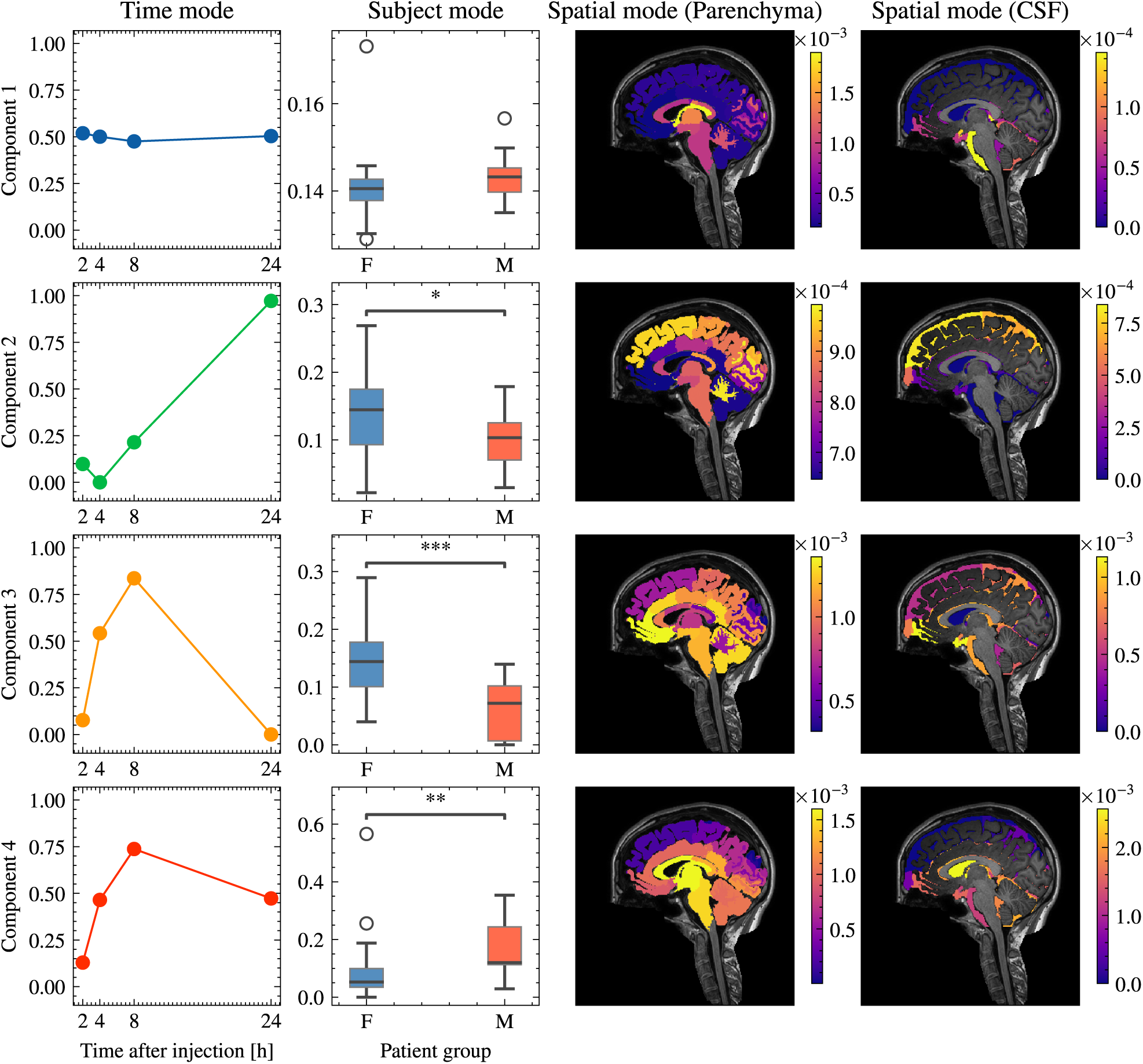
Time, subject and spatial modes obtained using a 4-component CP model on the REF subset of the dataset. We split the spatial component into parenchyma and CSF to view the dynamics in each region more clearly.

**Figure S8:**
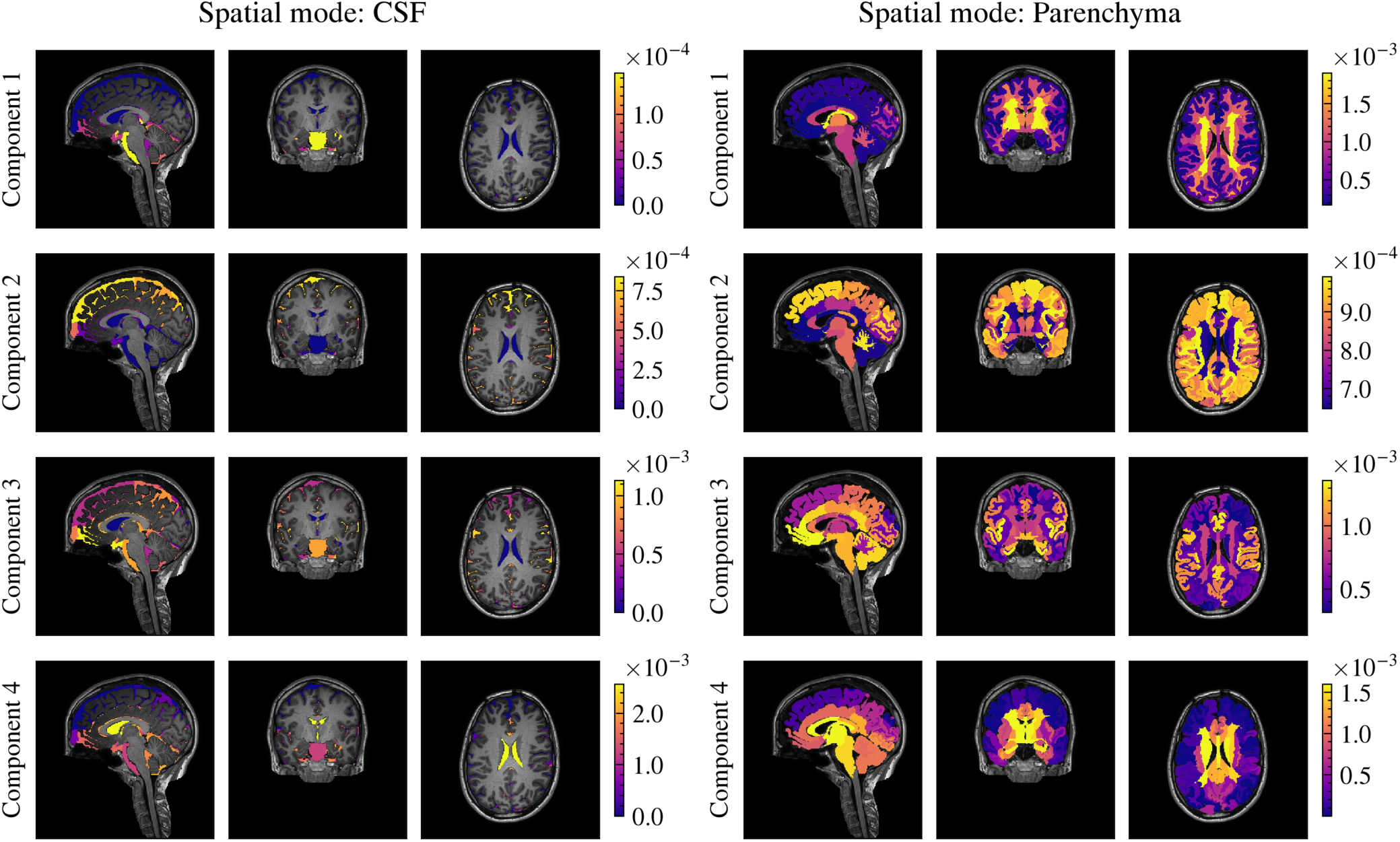
Sagittal, coronal and axial slices of the spatial mode for each component on the REF subset of the dataset. Provides a detailed spatial view of Figure S7.

**Figure S9:**
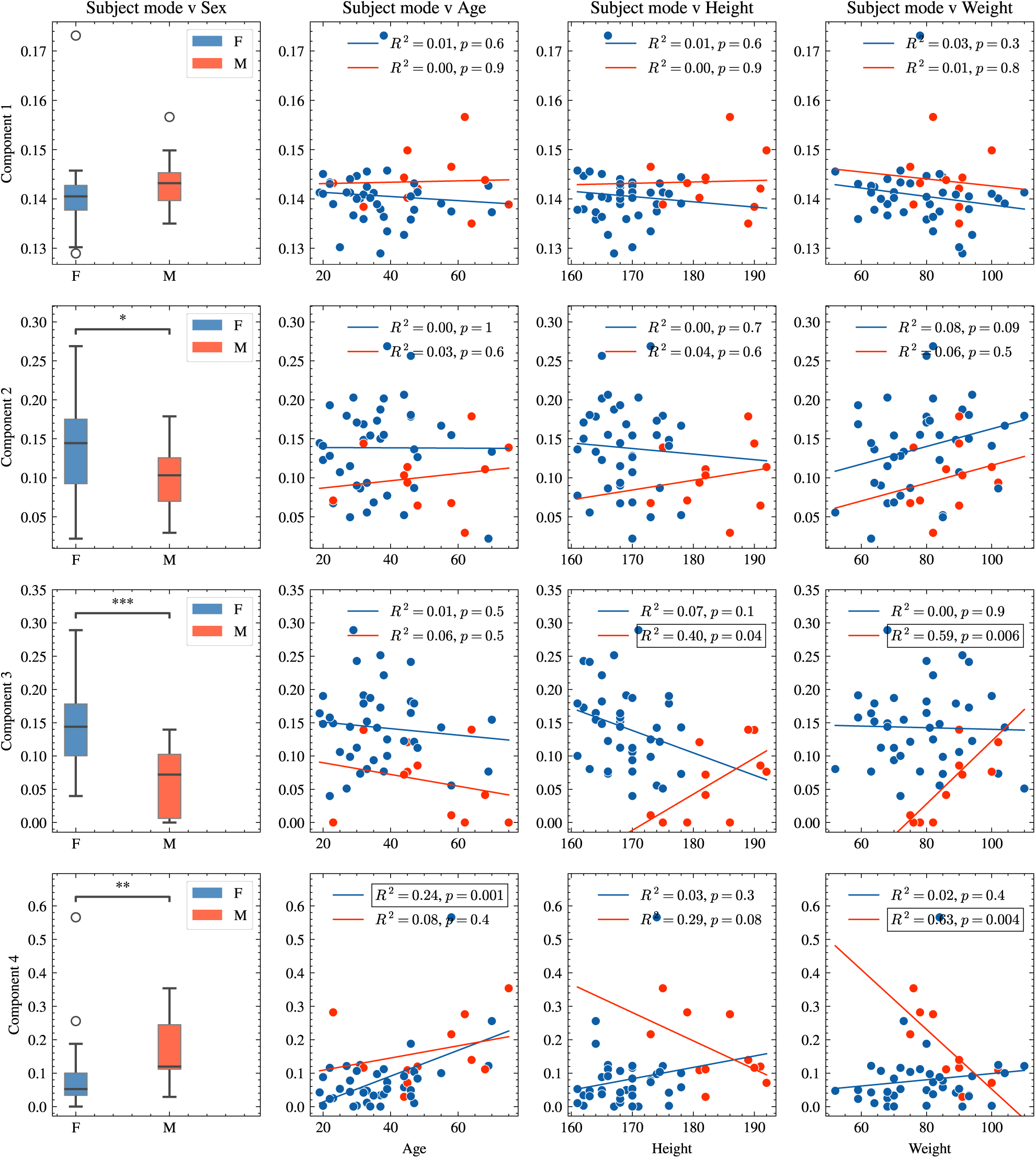
Subject mode coefficients obtained for each component as a function of various subject characteristics on the REF subset of the dataset. We compare the two groups (male and female), the third and fourth components are found to be significantly different with regard to sex (Mann-Whitney U-test, *p <* 0.05). The subject mode coefficient is also plotted with respect to age, height and weight for each of the two groups.

## Notes

### Competing Interest Statement

The authors have declared no competing interest.

